# Αlpha-Synuclein Induced Immune Response Triggers Parkinson’s Disease-Like Symptoms

**DOI:** 10.1101/2024.05.27.596130

**Authors:** Rebekah G. Parkinson, Tony Xu, Jacob Martin, Zizheng Xian, Ilvana Ziko, Jessica A. Pettitt, Alexandre RCom-H’Cheo-Forgues, Rebecca Buckland, Sarah L. Gordon, Christopher Parish, Anne Brüstle, Nathalie Dehorter

## Abstract

Increasing evidence suggests that Parkinson’s disease is an autoimmune disorder, with findings of elevated peripheral blood mononuclear cell in patients, and antigenic properties of α-synuclein driving both the innate and adaptive immunity. Yet, how the interaction of α-synuclein and a specific immune response participates to Parkinson’s disease ontogenesis has remained unanswered. Here, we reveal that autoimmune response to an α-synuclein antigen underlies Parkinson’s disease. We demonstrate that autoimmunity mediated by CD4+T cell activation with α-synuclein α-syn_61-75_ antigen is required to lead to immune cell infiltration and localized inflammation in the substantia nigra, triggering dopaminergic cell neurodegeneration and deficits in locomotion and gait kinematics. This study offers the first immune-induced mouse model that recapitulates all features of Parkinson’s disease to study the mechanisms triggering disease onset. It provides the basis for temporally tracking symptom development, exploring preventive strategies and prodromal therapeutic interventions in Parkinson’s Disease.

**In brief:** Peripheral α-synuclein immunization causes Parkinson’s disease-like symptoms in mice.

**Highlights:**

- Both CD4+ T cells and α-synuclein are essential for Parkinson’s disease ontogenesis.
- Peripheral injection of α-syn_61-75_ induces significant CD4+ T cell infiltration in the mouse brain.
- α-syn_61-75_ immunization is associated with inflammation, α-synuclein aggregation and dopaminergic cell loss in the substantia nigra pars compacta.
- Levodopa-sensitive motor symptoms are detected 8 weeks following α-syn_61-75_ immunization in mice.
- This study offers a novel autoimmune α-synuclein induced mouse model of Parkinson’s disease.

## INTRODUCTION

Parkinson’s disease (PD) is a progressive neurodegenerative disorder characterized by the selective loss of dopaminergic neurons in the substantia nigra (SN), leading to the cardinal motor symptoms of tremor, rigidity, akinesia, and bradykinesia (1). Although the aetiology of PD remains multifactorial and complex, emerging evidence suggests a pivotal role for the interplay between α-synuclein aggregation and the immune system. α-synuclein, a small soluble protein predominantly localized at presynaptic terminals demonstrates prion-like propagation and is posited to trigger immune-mediated mechanisms within both the peripheral and central nervous system (2). When aggregated, α-synuclein accumulates into amyloid fibrils (3) aiding in the degeneration of dopaminergic cells within the nigrostriatal pathway (4). Evidence from seed amplification assays reports that α-synuclein represents a robust Parkinson’s progression marker in humans providing high sensitivity to participants with PD and detection of prodromal cases (5). Research in recent years has also described the highly immunogenic nature of α-synuclein, identifying specific α-synuclein-derived epitopes that drive both innate and adaptive immunity, even in cases that precede the development of PD symptoms (6), suggesting that α-synuclein could participate to the early, prodromal stages of the disease. In this context, α-synuclein-specific CD4+ and CD8+ T cells have been found in the brains of people with PD, further supporting that α-synuclein may act as an autoantigen, triggering adaptive immune responses (7).

T cells have been implicated in neuroinflammation and with their activation, release of pro-inflammatory cytokines such as interleukin (IL)-17, interferon gamma (IFNΓ-γ), and Granulocyte-Macrophage–colony stimulating factor (GM-CSF), further exacerbating neuronal damage (8). A recent study by Garretti et al. (9) also reported specific interaction between the immune CD4-positive T cells (CD4+) and α-synuclein_32-46_ (α-syn_32-46_) peptide, associated with common genetic variation in the HLA region and correlated with late-onset sporadic PD (6), in leading to enteric features of prodromal PD in mice. However, immunization to α-syn_32-46_ failed to trigger neurological and motor symptoms of PD. Therefore, determining the critical association of α-synuclein autoimmunity in prodromal idiopathic PD, which represents more than 80% of the known PD cases, remained to be determined.

Here, we aimed to uncover the triggers of stereotypical Parkinson’s symptoms, including dopaminergic cell loss, α-synuclein aggregation, inflammation in the substantia nigra pars compacta (SNc), and related motor deficits. We speculated that the specific α-synuclein peptide (α-syn_61-75_) associated with immune CD4+ T cells in idiopathic patients (7) could underlie PD development. This work reports the first immune-induced model recapitulating Parkinson-like symptoms in mice. It represents a key tool to study the prodromal mechanisms underlying the ontogenesis of the disease.

## RESULTS

### α-synuclein immunization elevates CD4+ T cell levels in mouse

To decipher the associations between the CD4+ T helper cells and α-synuclein in the ontogenesis of idiopathic PD, a systemic immune reaction against α-synuclein was used by subcutaneously injecting α-syn_61-75_ (**Fig. 1A, B**) or the whole α-synuclein protein (**Fig. S1A, B**) in Complete Freud’s Adjuvants (CFA) to trigger an α-synuclein specific response. Blood immunophenotyping and cell counting performed one week before (*i.e.* baseline) and one week after induction demonstrated both sham and α-synuclein immunized mice with a robust inflammatory response in the blood, identified by marked elevation of white blood cells (WBCP), monocytes, and neutrophils, compared to non-injected mice (**Fig. 1C; S1C)**. The immune activation was similar for either human or mouse α-synuclein (**Fig. S1D, E**), with no modifications to standard blood measures unrelated to inflammation such as red blood cells **(Fig. S1F)**, haematocrits **(Fig. S1G)**, and haemoglobin **(Fig. S1H)**. We also observed that the transient immune response was maintained till 8 weeks (**Fig. S1I-K**), indicating sustained activation of the immune system following single α-synuclein immunization. Given the strong link of α-synuclein to CD4+ T helper cells (6), we first assessed sustained activation of Th cells indicated by their expression of the activation marker CD44 at 8 weeks post immunization (wpi). We performed flow cytometry staining (Gating strategy in Methods and **Fig. S2A**), finding mice immunized with α-syn_61-75_ displayed a significant increase of activated CD4+ T cells compared to sham induced mice indicating some α-syn_61-75_specific activation (**Fig. 1D**). To investigate the nature of this specific response, we isolated cells from cervical lymph nodes (cLN) at an earlier timepoint (*i.e.* 3 wpi) and stimulated them ex vivo with α-syn_61-75._ Following intracellular cytokine restimulation **(Fig. S2B**), we revealed an increase of T cells producing IL-17 and GM-CSF (**Fig. S2C, D**), cytokines strongly linked to neuroinflammation in general (10), as seen for example in the experimental autoimmune encephalomyelitis (EAE) model of neuroinflammation (11). To further explore the specific effect α-synuclein has on T cell populations, we investigated potential immunological memory and affinity after initial exposure and priming to α-synuclein (*i.e.* two weeks after induction), utilizing *in vitro* lymphocyte cultures stimulated in α-synuclein protein or α-syn_61-75_. Interestingly, α-synuclein immunized lymphocytes did not demonstrate any significant increase in CD4+ T cells activation when restimulated with the same protein (**Fig. S2C)**. However, restimulation with α-syn_61-75_ showed increased activation of proinflammatory cytokines (**Fig. S2C)**. This elevation was greatest in GM-CSF+ IL-17+ populations in mice that were immunized to α-syn_61-75_ and restimulated in the same peptide, indicating specific immunological memory to α-syn_61-75._ Since these results indicated significant effects of the α-synuclein immunization on the adaptive immune system, we investigated further their involvement in the specific immune response to α-synuclein. Using random forest selection methods via Boruta, we found increased populations of GM-CSF+, IFN**γ**+ and IL-17+ producing cells in α-synuclein immunized mice compared to sham conditions, at 8 weeks post-induction (**Fig. 1E-G**). This suggests that the increase in GM-CSF+, IFN**γ**+ and IL-17+ cells persist beyond the transient immune activation triggered by CFA-MT administration alone. Interestingly, GM-CSF+ and IL-17+ producing cells are produced by pathogenic Th17+ cells which are strongly linked to neuroinflammation, as seen in the experimental autoimmune encephalomyelitis model (EAE; (12) (13). Th17 cells and the innate neutrophil granulocytes recruited through Th17 cells are the hallmark cells of type III immune responses (14) and indeed, we still observed an increase of neutrophil granulocytes (CD11b+Ly6G+ cells) at week 7 post-induction (**Fig. 1 H, I)**. These results indicate persistence of CD4+ mediated type III response.

**Figure 0.** Immunization with α-synuclein triggers specific immune response which leads to Parkinson’s like alterations. Type 3 immune response to α-synuclein involving CD4+ T cells stimulate IFNy, IL-17, and GM-CSF which triggers neutrophils, induces dopaminergic cell death, inflammation and motor deficits in male and female C57bl/6 mice, mimicking idiopathic Parkinson’s disease.

**Figure 1.**
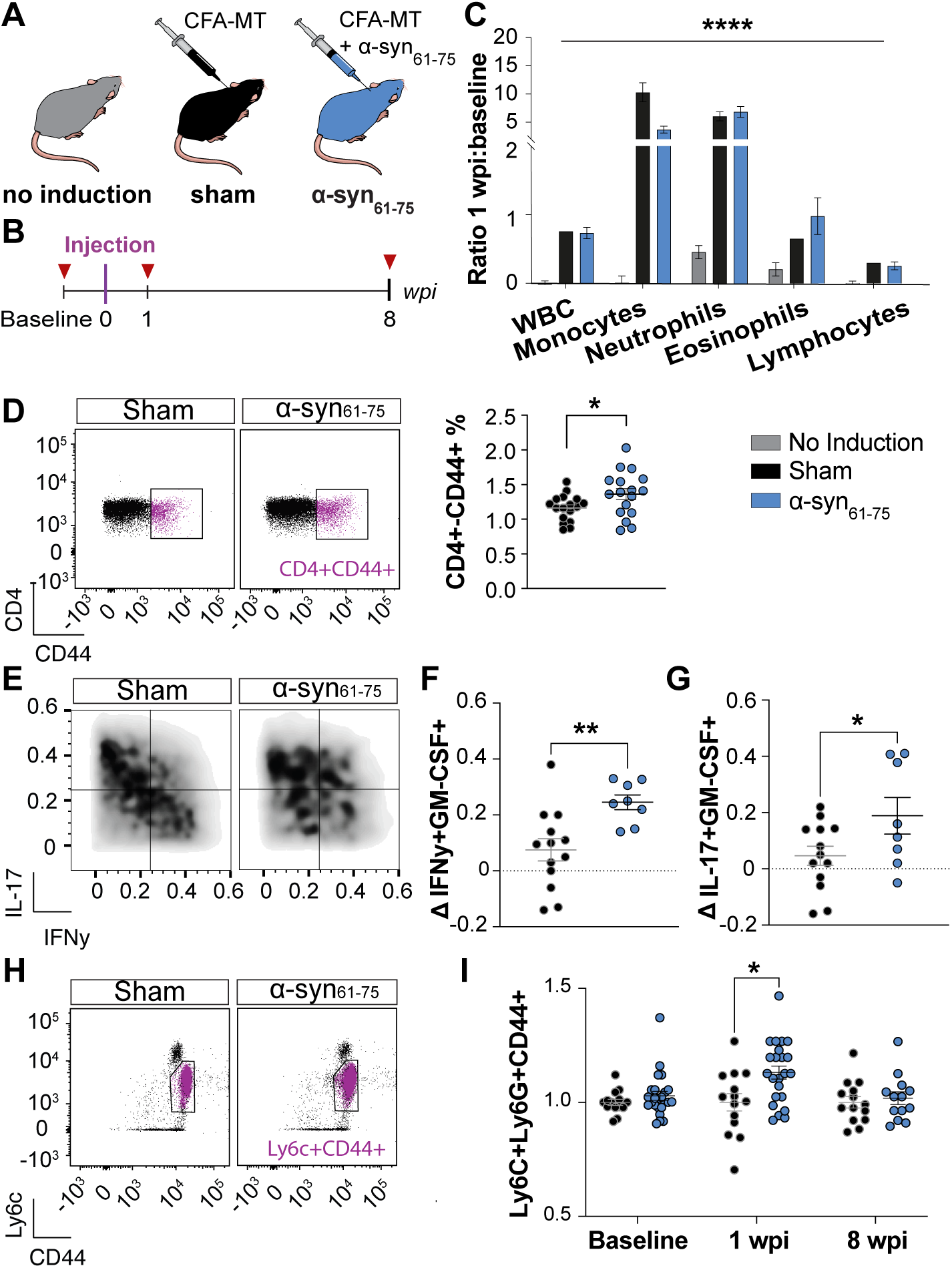
Immunization to α-synuclein triggers robust and persistent CD4+ T cell activation. (A) Experimental groups. (B) Timeline including blood collection (red triangle), injection (purple), and cervical lymph node (cLN) extraction. Week post immunization: wpi. (C) Full blood counts of white blood cells (WBC), monocytes, neutrophils, eosinophils, and lymphocytes quantified as a ratio of 1 wpi over baseline. Two-way ANOVA condition effect F(2,280) = 18.08, p < 0.0001. No-induction (n = 10), sham (n = 30), and α-syn_61-75_ immunized (n = 19) mice. (D) Left: Representative flow cytometry plots of CD4 and CD44 (double positive in blue) in blood, pregated according to Fig. S2. Right: CD4+CD44+ cells as a percentage of live cells for sham (n = 17) and α-syn_61-75_ immunized (n = 17) mice, p = 0.0355. (E) Cells isolated from cLN on 2 wpi were cultured ex vivo with and without α-syn_61-75_ for 4h, before intracellular cytokine staining. Shown is the % of GM-CSF+ (left) and IL-17+ (right) cells as the difference between cells stimulated with α-syn_61-75_ minus the cells without α-syn_61-75_ stimulation as an indicator for α-syn_61-75_-unspecific of background cytokine production. One-way ANOVA F(X) = X, p < X. Tukey multiple comparisons sham vs α-syn_61-75._ (F) Density plots of IL-17 and IFNy expression within GMCSF+ blood populations at 8 wpi normalized to the baseline expression. (G) Mean fluorescent intensity (MFI) of IFNy+ GM-CSF+ (left, p = 0.0059) and IL-17+ GM-CSF (right, p = 0.0457) cells, at 8 wpi relative to the baseline. Sham (n = 13), α-syn_61-75_ immunized (n = 8) mice. (I) Representative flow cytometry plots of CD11b and Cd11c in blood, pregated according to Fig.S2, and overlayed with Ly6c+CD44+ cells (blue). (J) MFI of Ly6g+CD44+ pregated from CD11b+Ly6c+ for sham (n = 14) and α-syn_61-75_ immunized (n = 13-24) mice. Mixed-effects condition effect p = 0.0097. Tukey’s multiple comparisons sham vs α-syn_61-75_ at 1 wpi, p = 0.0107. SData points represent individual mice values. Data is shown as mean ± SEM. Significance represented with p < 0.05: *; p < 0.01: **; p < 0.001: ***, p < 0.0001: ****. Non-paired Student T test otherwise stated. See also Fig. S1&2.

### Immunization of α-syn61-75 induces Parkinson-like neurological and behavioural alterations in mice by CD4^+^ T cell-mediated infiltration

We hypothesized that the shift in neutrophils and T cell populations (**Fig. 1**) could be due to immune cell infiltration from blood to brain. We therefore monitored adaptive immune cell infiltration two weeks post-immunization and directly quantified CD4+ T cells in the mouse brain lateral ventricles. We found significant CD4+ T cell infiltration in α-syn_61-75_ immunized mice, compared to sham mice (**Fig.2A, B**). We monitored pathogenic infiltration and did not find any difference in CD3+ (*i.e.*, CD4+ and CD8+ cells), CD4+ and Ly6b+ (corresponding to neutrophils and activated monocytes) cell densities in the SNc of α-syn_61-75_ immunized mice 8 weeks post-induction (**Fig. S3A-C**).

**Figure 2.**
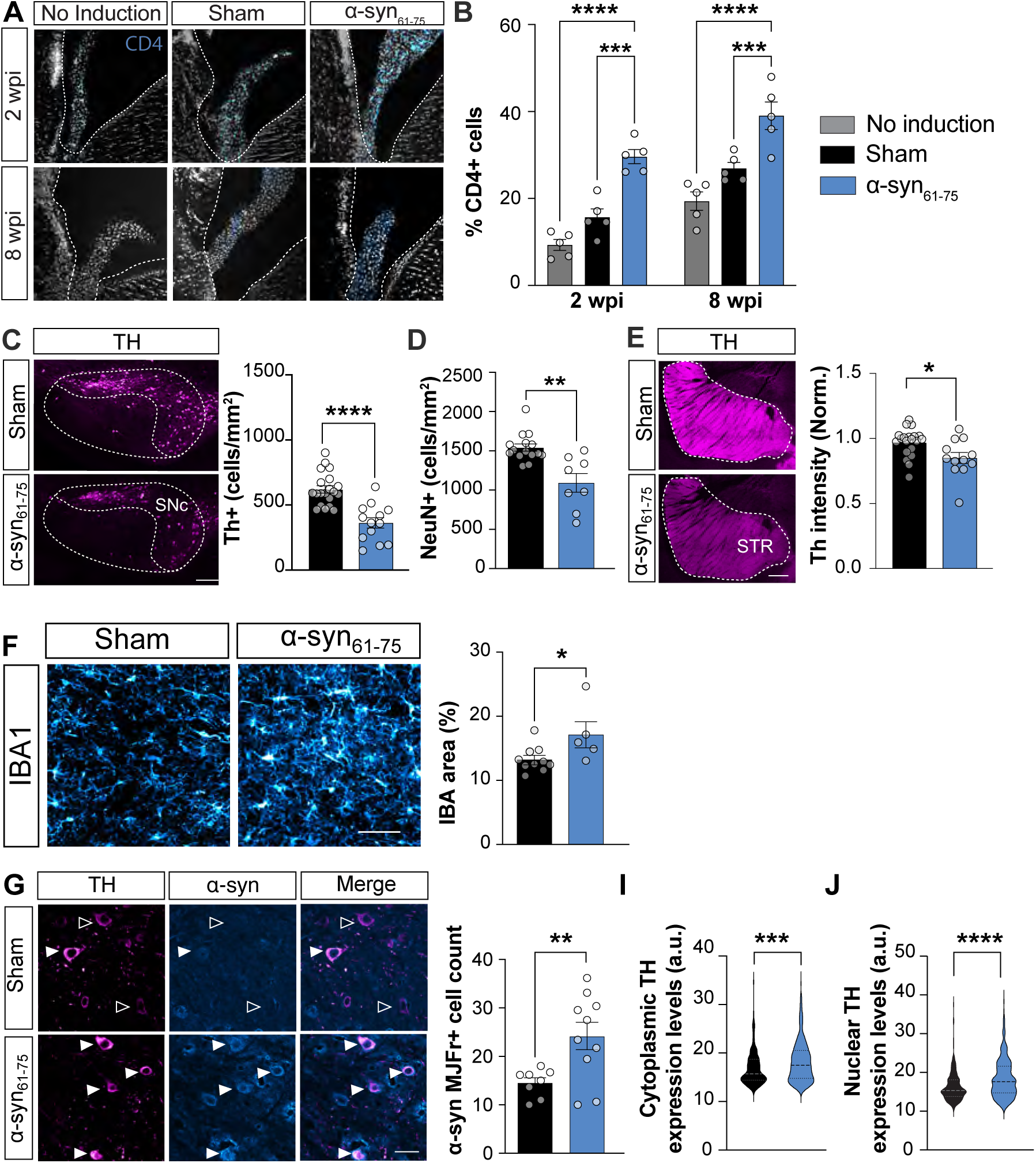
Cellular alterations in the brain of mice immunized to α-syn_61-75_. (A) Representative images of CD4+ T cell infiltration in the choroid plexus of the lateral ventricle of α-syn_61-75_ immunized mice (n = 5) compared to control (no-induction) and sham mice (n = 5) at 2- and 8-weeks post injection (wpi). Scale = 50 µm. (B) Two-way interaction effect p < 0.0001. Tukey’s multiple comparisons of α-syn_61-75_ immunized mice to non-induced (p < 0.0001) and sham (p = 0.0002) mice at 2 wpi, and α-syn_61-75_ immunized mice to non-induced (p < 0.0001) and sham (p = 0.0008) mice at 8 wpi. See also Fig. S3. (C) Left: representative images of tyrosine hydroxylase (TH+) dopaminergic cells in the substantia nigra pars compacta (SNc). Scale = 100 µm. Right: TH+ dopaminergic cell density in sham (n = 20) and α-syn61-75 injected mice (n = 13) at 8 weeks post injection, p < 0.0001. (D) SNc NeuN density in sham (n = 15) and α-syn61-75 injected mice (n = 8) at 8 wpi, p = 0.0011, Mann-Whitney test. (E) Left: representative images of TH+ dopaminergic afferences in the striatµm (STR). Scale = 500 µm. Right: TH expression levels in the striatµm normalized to sham. Sham (n = 20) and α-syn_61-75_ injected mice (n = 12) at 8 wpi, p = 0.0132. (F) Left: representative images of IBA1+ microglia neuropil; Scale = 50 µm. Right: IBA1+ percentage area in the SN of sham (n = 10) and α-syn_61-75_ immunized mice (n = 5) at 8 wpi, , p = 0.0280, Mann-Whitney test. (G) Left: representative images of MJFR-14-6-4-2+ colocalized with TH+ cells in the SNc (filled arrow). Scale = 30 µm. Increase in α-synuclein aggregates in α-syn_61-75_ immunized mice (n = 10) compared to sham mice (n = 8); p = 0.0081, T-test with Welch correction. Fluorescent intensity of MJFR-14-6-4-2 inside the dopaminergic cell cytoplasm (H; p < 0.001, Mann-Whitney test) and nucleus (I; p < 0.001, Mann-Whitney test; 199 sham cells., 334 α-syn_61-75_ immunized cells). Data points represent individual mice values. Data is shown as mean ± SEM. Significance represented with p < 0.05: *; p < 0.01: **; p < 0.001: ***, p < 0.0001: ****. Non-paired Student T test otherwise stated.

We next aimed to verify the neurological consequences of immune cell infiltration in the mouse brain. Since the hallmark of PD is characterized by dopaminergic cell neurodegeneration, we quantified the density of tyrosine hydroxylase (TH+) dopaminergic neurons within the SNc. Whilst non-induced and sham mice groups did not show any difference in TH+ dopaminergic cell density (**Fig. S3D,** we identified an average of 40% dopaminergic cell loss at 8 weeks in the α-synuclein protein **(Fig. S3E, F)** and α-syn_61-75_ injected mice **(Fig. 2C)** which persisted until 40 weeks (Data not shown) and which was associated with dopaminergic loss in the striatum **(Fig. 2E; S3G)**. No differences were found in TH+ dopaminergic cell density in other dopaminergic structures such as the ventral tegmental area (**Fig. S3H)**. Another key feature observed in Parkinson’s patients is localized inflammatory activity (2). As such, we found increased expression of IBA1+ microglia in the SN after 8 weeks in α-synuclein protein **(Fig. S3I)** and α-syn_61-75_ **(Fig. 3F)** immunized mice compared to the controls, whilst no neuroinflammation was detected in the striatum, cortex or hippocampus (n=5, 4 & 4 mice; a-synuclein vs sham: p=0.9943, 0.9950 & 0.9493, respectively; Two-way ANOVA, Tukey’s multiple comparison test; Data not shown). These results suggest the presence of a specific inflammatory response to α-synuclein within the SN. PD is also characterized by intraneuronal inclusions of α-synuclein within SNc dopaminergic neurons (15). Interestingly, we found increased number of TH+ dopaminergic cells that were colocalized with α-synuclein in mice immunized with the specific α-syn_61-75_-injected mice **(Fig. 2H-J)**.

**Figure 3.**
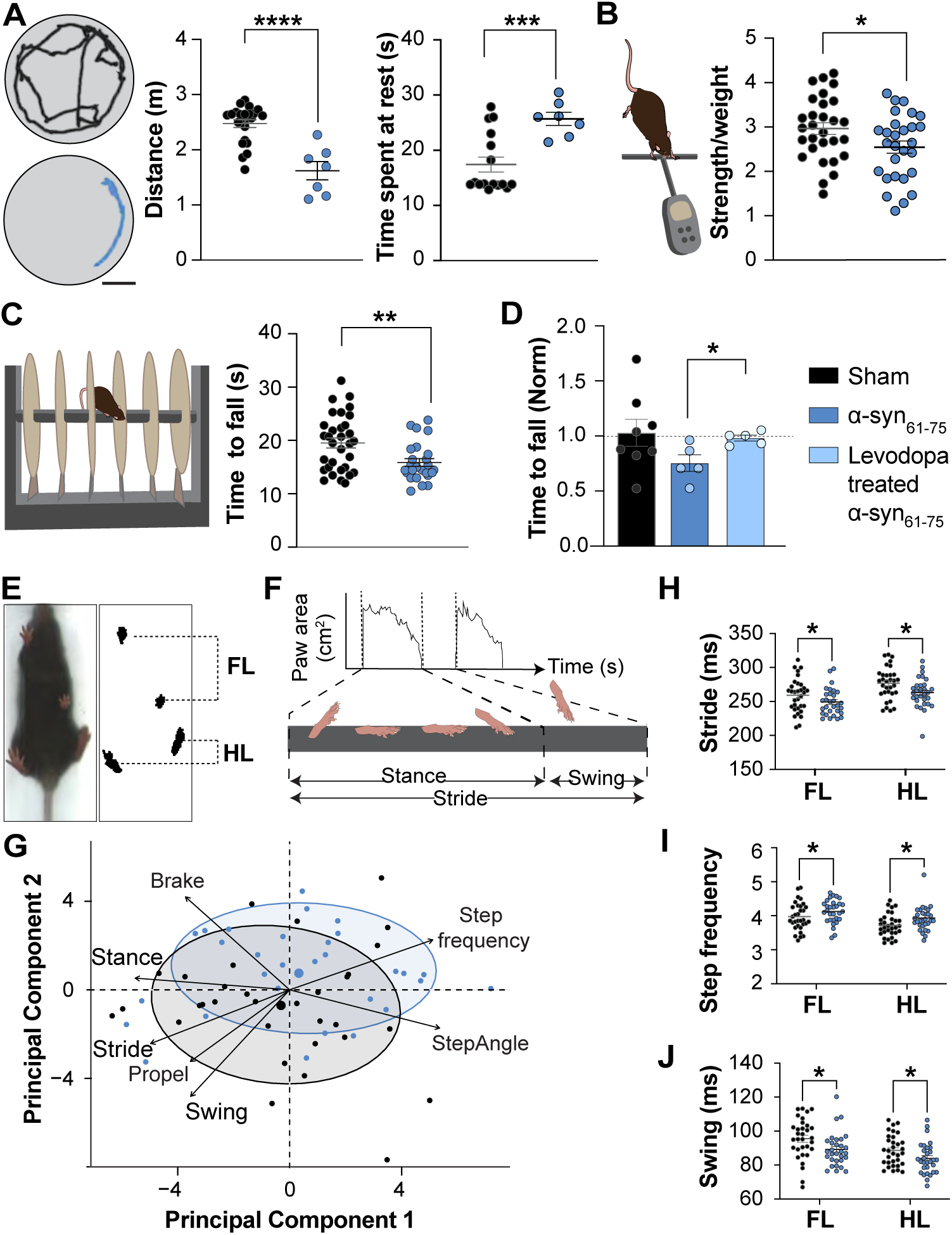
α-syn_61-75_ immunization triggers motor deficits in gross locomotion, coordination, and fine gait kinematics. (A) Left: Representative schematic of mouse locomotor tracking in open field arena from sham (top) and α-syn_61-75_ immunized mice (bottom). Distance travelled (middle, p = 0.0006) and time spent at rest (right, p = 0.0002, T-test with Welch correction) in arena at 8 weeks post-immunization (wpi) in sham (n = 16) and α-syn_61-75_ immunized mice (n = 7). (B) Grip strength at 8 wpi in sham (n = 29) and α-syn_61-75_ mice (n = 28), p = 0.0361. (C) Accelerating rotarod test at 8 wpi, sham (n = 32) α-syn_61-75_ immunized (n = 25) mice, p = 0.0029, T-test with Welch correction. (D) Rotarod test scores normalized to sham (n = 8) mice between α-syn_61-75_ immunized mice treated for 15 days with Levodopa (n = 5), compared to untreated α-syn_61-75_ immunized mice (n = 5); p = 0.0213. (E) Left: Screen capture of video footage displaying the mouse on the DigiGait apparatus. Middle: Mask area of each paw. Right: Illustration of some parameters analyzed: paw area, paw angle and midline distance. (F) Principal component analysis of 30 hindlimb DigiGait parameters clustered by condition at 8 wpi. Parameters shown with high variable loadings for principal component 2 (swing; 0.4600, brake; 0.3501, propel; 0.2080, stride; 0.1170, step frequency; 0.1013, stance; 0.0048, and step angle; 0.0044). See also Fig. S3J. (H-J) DigiGait metrics separated by forelimb (FL) and hindlimb (HL) from sham (n = 32) and α-syn_61-75_ immunized (n = 30) mice, at 8 wpi. (H) Stride duration. Two Way-ANOVA, condition effect F(1,121) = 8.461 p = 0.0043, Tukey’s multiple comparison p = 0.0022 vs sham. (I) Step frequency. Two Way-ANOVA, condition effect F(1,121) = 7.130, p = 0.0086, Tukey’s multiple comparison p = 0.0423 vs sham. (J) Swing duration. Two Way-ANOVA, condition effect F(1,121) = 9.564, p = 0.0025, Tukey’s multiple comparison p = 0.0130 vs sham. Data points represent individual mice values. Data is shown as mean ± SEM. Significance represented with p < 0.05: *; p < 0.01: **; p < 0.001: ***, p < 0.0001: ****. Non-paired Student T test otherwise stated. See also Fig. S4.

To investigate physical manifestations that could be triggered by these cellular deficits observed in α-synuclein immunized mice, we next performed a battery of behavioral tests. α-synuclein immunized mice did not demonstrate any accumulative weight increase post-induction and no differences in food consumption **(Fig. S4A, B**). In addition, no behavioral changes were observed between non-induced and sham mice at 8 weeks (**Fig. S4C, D)**, and between α-synuclein protein/α-syn_61-75_ immunized and sham mice at 2-, 4- and 6-weeks post-injection (**Fig. S4E, F)**. However, significant motor deficits were observed from 8 weeks post-induction with immunized α-synuclein mice travelling less distance and spending more time at rest in an open field arena, compared to sham mice **(Fig. 3A; Fig. S4G)**. In addition, mice presented weaker grip strength **(Fig. 3B; Fig. S4H)** and signs of motor deficits in the rotarod test **(Fig. 3C; Fig. S4I)**. Interestingly, 15 days of treatment with Levodopa, a dopamine replacement agent for the treatment of PD (16), restored motor alterations in α-syn_61-75_ injected mice **(Fig. 3D)**. To further assess motor alterations, we analyzed fine gait kinematics on a DigiGait treadmill **(Fig. 3E)**. α-syn_61-75_ immunized mice displayed various deficits in gait parameters compared to non-injected and sham control mice, including stride, stance or swing (**Fig. 3F-J; Fig. S4J)**. In addition, we found rest/sleep-related disturbances quantified by increased home cage activity (**Fig. S4K)**, which represent a behavioral feature observed in PD patients (17). Overall, these results confirm that α-synuclein immunization triggers Parkinson’s-like symptoms in wildtype mice.

### Parkinson’s like symptom ontogenesis depends on nascent α-synuclein and adaptive autoimmune activation

Since PD is a neurodegenerative disorder mostly affecting old populations (18), we next aimed to decipher whether age, which represents the largest risk factor for the development and progression of PD, could be an aggravating factor in disease ontogenesis in α-syn_61-75_ immunized mice. To test this, we investigated for any exacerbation of α-synuclein dependent immune activation, dopaminergic cell loss and related behavioral deficits in older mice (> 12 months), which displayed a higher number of neutrophils but the same number of lymphocytes and CD4+CD44+ cells, compared to young adult mice (**Fig. 4A**). It is also interesting to note that dopaminergic TH+ cell density and motor behavioral performance in rotarod in old mice were not different compared to the young adult mice (**Fig. 4B**). Following immunization with the α-syn_61-75_ (**Fig. 4C**), old mice recapitulated the alterations found in younger mice (**Fig. 2&3**), which included dopaminergic cell loss (**Fig. 4D**) and motor deficits (**Fig. 4E-G**). Interestingly, α-syn_61-75_ immunized old mice were also prone to refuse to run on the DigiGait apparatus, compared to sham mice (8 trials run out of 14, 7 mice vs 5 trials run out of 16, 8 mice, respectively; Chi-square test, p=0.15; Data not shown) and PCA analysis showed differences between both conditions in mice which ran on the DigiGait (n=5 sham and 3 α-syn_61-75_ injected mice; Student t test; p=0.03; Data not shown). However, we could not detect any worsening of the symptoms in α-syn_61-75_ peptide injected mice, compared to the sham group (TH+ cell density from 41% in young *vs* 36% loss in old mice), indicating that age does not increase the vulnerability to α-syn_61-75_ immune-dependent outcomes.

**Figure 4.**
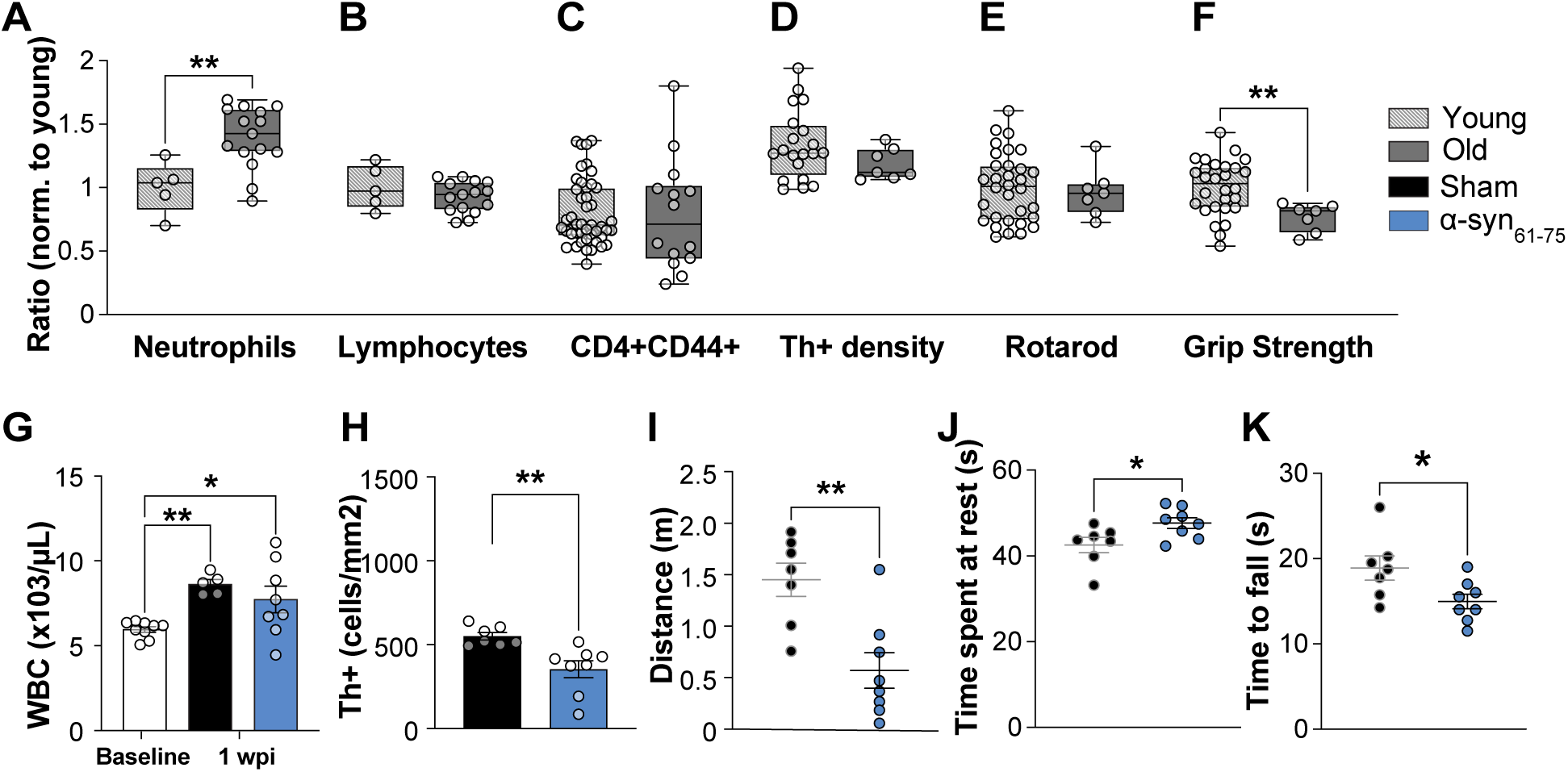
PD symptoms persist in old α-syn_61-75_ immunized mice. (A) Ratios of young (8-18 weeks) vs middle-aged (52 weeks) adult mice. Old (n = 15) mice had more whole blood counts of neutrophils at baseline compared to young (n = 5) mice, p = 0.0045. (B) No differences at baseline found in absolute lymphocytes between young (n = 5) and middle-aged mice (n = 15), p = 0.2955. (C) Flow cytometry of CD4+CD44+ % from young (n = 45) vs middle-aged (n = 14) mice, p = 0.8469, (D) SNc dopaminergic (tyrosine hydroxylase TH+) cell density (cells/mm^2^) of sham mice at 8 weeks post immunization (wpi), young (n = 20) and middle-aged (n = 7) mice, p = 0.1765. (E) Behavioral recordings of sham mice at 8 wpi, including time to fall off accelerating rotarod test (s), young (n = 32) and middle-aged (n = 7), p = 0.7189. (F) Significantly weaker grip strength in middle-aged (n = 7) compared to young (n = 28) mice, p = 0.0083. (G) Whole white blood cells (WBC) at baseline (n = 9), compared to 1 wpi from sham (n = 7) and α-syn_61-75_ immunized (n = 6) mice. One-way ANOVA, F(2,19) = 5.639, p = 0.0119; post-hoc Dunnett’s multiple comparisons versus baseline: sham p = 0.0238, α-syn_61-75_ p = 0.0176. (H) SNc dopaminergic cell density at 8 wpi sham (n = 7) and α-syn_61-75_ (n = 8) mice, p = 0.0047. (I) Distance travelled (m) in open field arena at 8 wpi in sham (n = 7) and α-syn_61-75_ immunized mice (n = 8), p = 0.0027. (J) Time spent at rest (s) in open field arena at 8 wpi in sham (n = 7) and α-syn_61-75_ immunized mice (n = 8), p = 0.0323. (K) Accelerating rotarod test at 8 wpi, sham (n = 7) and α-syn_61-75_ (n = 8) mice, p = 0.030. Data points represent individual mice values. Data is shown as mean ± SEM. Significance represented with p < 0.05: *; p < 0.01: **; p < 0.001: ***, p < 0.0001: ****. Non-paired Student T test otherwise stated.

We next investigated whether the injected α-syn_61-75_ alone was sufficient for the local reactivation of T cells in the central nervous system or whether the nascent α-synuclein protein was required to trigger the development of Parkinson’s like symptoms. To address that, we performed immunization of α-synuclein knockout (*α-syn^-/-^*) mice which are devoid of α-synuclein (19) (**Fig.5A**). As previously observed in wildtype mice, we observed significant increase in neutrophils (**Fig. 5B**) and WBC (**Fig. S5A**) in both sham and α-syn_61-75_ injected *α-syn^-/-^* mice, one-week post-immunization. Non-injected and sham *α-syn^-/-^* mice also presented with similar TH+ cell density and behaviour (**Fig S5B-D)**. However, contrary to the wildtype mice which display cellular and behavioural alterations after immunization (**Fig. 2&3**), we did not detect any dopaminergic TH+ cell loss (**Fig.5C**) and no behavioural changes between the sham and α-syn_61-75_ groups after 8 weeks (**Fig.5D**). This result indicates that nascent α-synuclein protein is required to trigger PD-like symptoms in mice following α-syn_61-75_ immunization.

**Figure 5.**
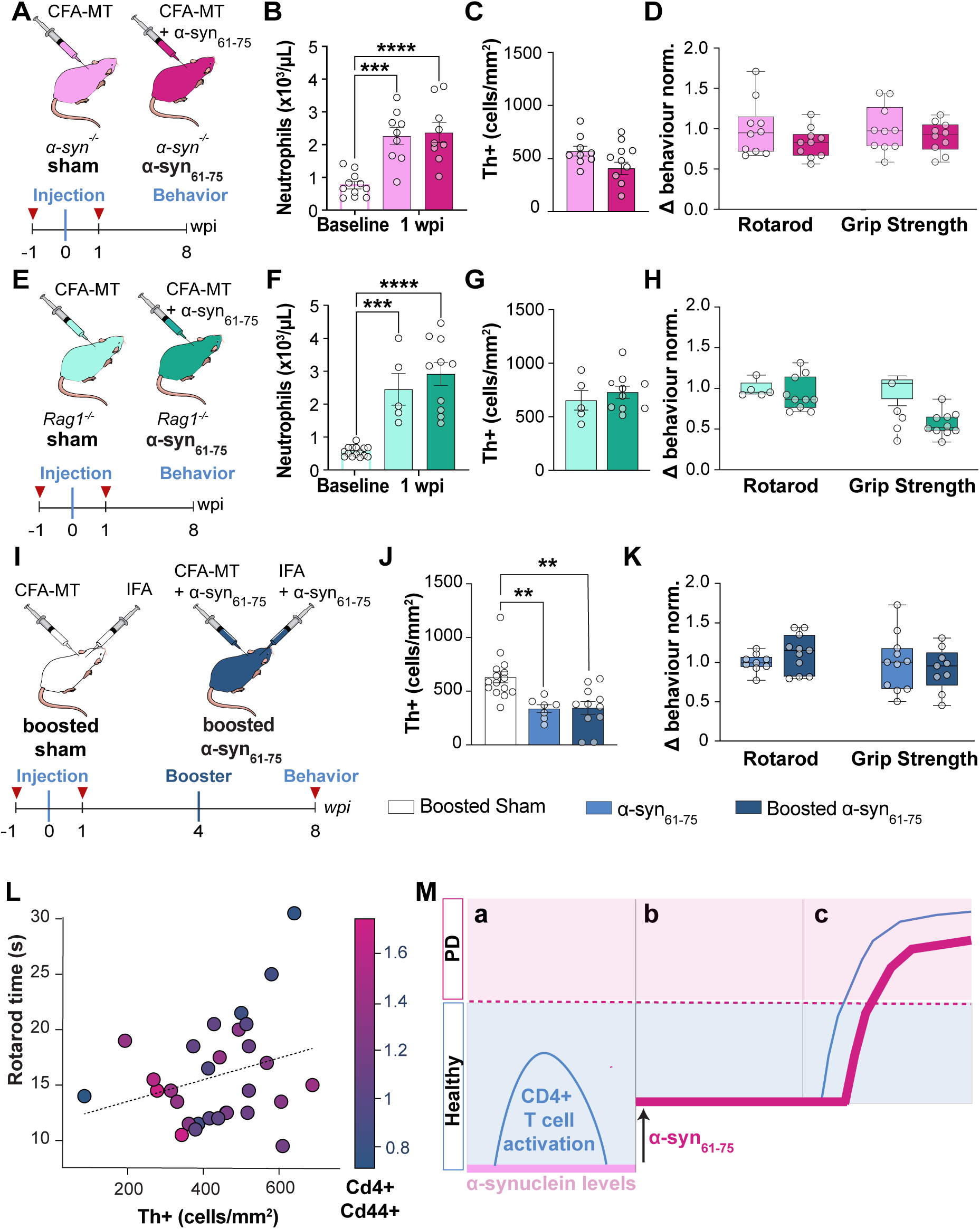
Cooperation of both immune T cell activation and α-syn_61-75_ is required to trigger PD symptoms. (A) Experimental groups for α-syn-/- mice and timeline including blood collection (red triangle), injection (blue, week 0) and behavioural assessment; wpi: week post-immunization. (B) Absolute neutrophils in whole blood at baseline (n = 11), compared to 1 wpi α-syn-/- sham (n = 9) and α-syn-/- α-syn_61-75_ immunized (n = 9) mice. One-way ANOVA test F(2,26) = 16.18, p < 0.0001. Dunnett’s multiple comparisons versus baseline: sham p = 0.0002, α-syn61-75 p < 0.0001. (C) No change in dopaminergic Th+ cell density in the SNc of α-syn_61-75_ immunized α-syn^-/-^ mice (n = 10), compared to sham α-syn^-/-^ mice (n = 10) at 8 wpi, p = 0.1865. (D) No change in motor behavior normalized to sham, between α-syn_61-75_ immunized α-syn^-/-^ mice (n = 10) and sham (n = 10) α-syn^-/-^ mice in rotarod test (p = 0.1625) and grip strength (p = 0.3772) at 8 wpi normalized to α-syn^-/-^ sham mice. (E) Experimental groups for Rag1^-/-^ mice, and timeline. (F) Absolute neutrophils in whole blood at baseline (n = 15), compared to 1-week post-immunization (wpi) sham (n = 5) and α-syn_61-75_ immunized (n = 10) mice. One-way ANOVA test F(2,27) = 31.50, p < 0.0001. Dunnett’s multiple comparisons versus baseline: sham p = 0.0001, α-syn_61-75_ p < 0.0001. (G) No change in dopaminergic Th+ cell density in the SNc of α-syn_61-75_ immunized Rag1^-/-^ mice (n = 10), compared to sham Rag1^-/-^ mice (n = 5) at 8 wpi, p = 0.8928. (H) No change in motor behavior normalized to sham, between α-syn_61-75_ immunized Rag1^-/-^ mice (n = 10) and sham (n = 5) Rag1^-/^ mice in rotarod test (p = 0.5967) and grip strength (p = 0.9283) at 8 wpi normalized to Rag1^-/-^ sham mice. (I) Experimental groups and timeline for mice receiving boosted immunizations at 4 wpi (dark blue). (J) SNc Th cell density of boosted sham (n = 15), compared to single injected α-syn_61-75_ immunized mice (n = 7), and boosted α-syn_61-75_ immunized (n = 11) mice. One-Way ANOVA test F() = X, p = X. Dunnett’s multiple comparisons versus boosted sham: α-syn_61-75_ p = X, and boosted α-syn_61-75_ p = X. (K) No change in motor behavior between α-syn_61-75_ immunized boosted mice (n = 11) and single injected α-syn_61-75_ immunized mice (n = 9). Rotarod test (p = 0.1894) and grip strength (p = 0.5834) at 8 wpi normalized to single injection of α-syn_61-75_ mice. (L) Correlation of rotarod test performance, is positively correlated with SNc Th+ cell density at 8 wpi (n = 29, p = 0.022, linear model regression test). These parameters are negatively correlated with percentage of CD4+CD44+ in live blood cells (p = 0.006, linear model regression test). (M) Schematic representation of the posited interplay between α-synuclein and CD4+ immune activation in PD(a) In normal conditions (*i.e.* healthy, blue area), CD4+ T cell activation (blue line) is not sufficient to trigger PD (disease threshold: dotted line). (b) Elevated levels of α-synuclein alone (*i.e.* without B/T cell immune activation) does not induce PD-like symptoms. (c) High levels of α-synuclein, combined with specific α-synuclein-dependent immune CD4 activation trigger PD-like symptoms in mice (dark pink area). Data points represent individual mice values. Data is shown as mean ± SEM. Significance represented with p < 0.05: *; p < 0.01: **; p < 0.001: ***, p < 0.0001: ****. Non-paired Student T test otherwise stated. See also Fig. S5.

Since the autoimmune activation triggers the development of Parkinson’s like symptoms in wildtype mice primed to α-synuclein, we next aimed to confirm a causative link between lymphocytes and the onset of PD symptoms. We performed α-syn_61-75_ immunization in immune-deficient *Rag1^-/-^* mice that lack B and T lymphocytes (20) **(Fig. 5E; Fig. S5E-G)**. Whilst both sham and *Rag1^-/-^* mice injected with α-syn_61-75_ displayed elevated neutrophils a week post α-syn_61-75_ injection indicating successful immunization **(Fig. 5F)**, they did not display any TH+ dopaminergic cell loss in the SNc **(Fig. 5G)** and no behavioural alterations at 8 weeks post immunization **(Fig. 5H)**. These results demonstrate that lymphocytes are essential for triggering Parkinson’s-like symptoms in mice following α-syn_61-75_ immunization.

We next posited that further enhancing the immune system could boost the reaction against α-synuclein immunization and ultimately could intensify disease course and exacerbate symptoms. Mice initially immunized to α-syn_61-75_ received an additional booster injection 4 weeks post induction with a-syn in incomplete Freud’s adjuvance (**Fig.5I)**. As expected, we found that the booster injection produced a significant elevation of white blood cell counts compared to the baseline, and to the single injected α-synuclein (**Fig. S5H**) or α-syn_61-75_ immunized mice **(Fig. 5F)**. However, despite this additional immune response, we did not observe any further increase in neutrophils (**Fig. S5I**) and CD4+ T cells (**Fig. S5J**) post-booster injection, compared to α-syn_61-75_ immunized mice. Most importantly, we found no evidence of worsened dopaminergic cell degeneration **(Fig. 5G),** or motor deficits in the α-syn_61-75_ boosted mice compared to single α-syn_61-75_ injected mice **(Fig. 5I)**. The lack of changes between boosted and single-immunized mice suggests that booster immunization does not enhance Parkinson’s like symptoms in α-syn_61-75_ immunized mice. These results also indicate that there could be a ceiling effect in the CD4-dependent α-synuclein recruitment. Significant positive correlation between elevated CD4+CD44+ T cells, dopaminergic cell loss, and poor rotarod motor performance (**Fig. 5M**) rather suggests a complex interplay.

Overall, these results reveal that both nascent α-synuclein protein and B/T cells are individually required after α-syn_61-75_ priming to trigger PD-like symptoms in mice. They also demonstrate a ceiling effect of the immune-dependent PD induction, with no effect of immune boost and no age-related enhancement of the immune response or worsening of immune α-syn_61-75_-induced symptoms.

## DISCUSSION

It has been well established that the immune system plays an important role in PD pathophysiology (2). Here, we questioned whether and how the immune system and α-synuclein interact during PD onset. We show that peripheral immunization against α-synuclein leads to PD-like symptoms in wildtype mice, including dopaminergic cell loss, inflammation and α-synuclein aggregation in the SNc, and motor impairments.

Whilst subtle changes in immune cell response were detected compared to the MS model (11), we still observed a type 3 immune response. We reveal a peptide-specific elevation of inflammatory CD4+ T cell cytokines including GMCSF, IL-17 and IFNy (**Fig.1**) and in particular, the double positive IFNy+IL-17+ cells, which represent pathological immune cells containing a full arsenal of cytokines. As there was also an increase in activated Ly6G+ and Ly6C+ populations in α-syn_61-75_ injected mice one-week post-induction, we suspect that the innate immunity, specifically polymorphonuclear cells, plays a key role at the initial stage of disease course and leads to peripheral CD4+ T cell activation and infiltration in the brain.

Furthermore, by utilizing the *Rag1^-/-^* model incapable of mounting an adaptive immune response due to the lack of B and T cells (20) **(Fig. 5E; Fig. S5E-G)**, we could further demonstrate that this immune induction relies on the adaptive arm of the immune system (**Fig. 5 E-H**). We confirm the crucial role of CD4+ T cells in triggering PD-like features, finding that the absence of CD4+/CD8+ T cells in *Rag1^-/-^*deficient mice prevents the appearance of symptoms in α-synuclein immunized mice. It is important to note that the presence of endogenous α-synuclein is also required to induce PD-like pathology, as *a-syn^-/-^* mice did not display PD-like symptoms when immunized with α-syn_61-75_. This could be because the autoimmune response is specifically triggered by endogenous α-synuclein in the body, or the increased amount of a α-synuclein available for antigen presentation is crucial during the induction process.

Interestingly, in contrast to other immunization-induced neurodegenerative models such as the Experimental Autoimmune Encephalomyelitis (EAE) model (20), changes in IL-17, GM-CSF and IFNγ were subtle and worked in combination to trigger the observed phenotype. In addition, we did not observe a single immune cell type involved in the mechanism of induction, but rather several of them. Thus, we propose to consider a holistic view involving both T cells and neutrophils to explain the onset of PD, with disease susceptibility increasing with higher endogenous levels of α-synuclein, elevated CD4+ T cells and neutrophils. We further suggest that the immune risk factor is critical to PD ontogenesis. Whilst the booster immunization was not effective in enhancing a significant immune response to α-synuclein, nor worsening symptoms in our conditions (**Fig. 5**), we appreciate that assessing substantial alterations following immune activation 4-week post-booster injection could have been too early and that a booster injection at 4 weeks post-immunization may have been too late (*i.e.*, when immune cells are already primed and infiltrated). More investigation on the timing and impact of the immune activation in PD ontogenesis is therefore required. Interestingly though, despite the booster immunization effectively elevating whole white blood cell counts, it did not specifically increase neutrophil or CD4+ activity. This may also explain why symptoms were not worsened.

As Parkinson’s disease is an age-related disorder, this study investigated whether age aggravates α-synuclein dependent immunization. Corresponding to the theory of inflammation (*i.e.* age-related increase in the peripheral dormant levels of pro-inflammatory markers (21), we observed increased neutrophils in middle-aged (12-month-old) mice, compared to young mice (**Fig.4**). However, we did not detect worsening of PD-like symptoms when these mice were injected with α-syn_61-75_. While age may contribute, the precise timing of α-synuclein immunization or combination of events over one’s lifetime may be critical in disease onset. Future investigations should aim at manipulating the temporal stage of immunization and immunizing aged mice (> 2 years old).

This study suggests that combination of an immune reaction and high basal levels of α-synuclein increases susceptibility to PD and agrees with current clinical research that investigates this association, in the context of recent pandemics (22). This work is also in line with a recent article by Garretti et al who reported prodromal features of genetic PD in the gut, with an altered Th1+/Th17+ profile and some altered associated gene profiles in PBS CFA vs CFA α-synuclein immunized groups (9). However, in contrast to our data the authors did not observe clear functional differences of their T cell responses since the analysis was not verified with intracellular staining or cytokines stimulation to go beyond correlation between α-synuclein and the altered profile. This discrepancy could be explained by different environmental elements and microbiome, as described for other immunization-based mouse models such as the EAE model of MOG 35-55 inducing MS, where a threshold is required to induce a robust immune response (11). We performed the experiments in two different laboratories and three different facilities, adjusting levels of pathogens, ensuring reproducibility of the phenotype under different conditions. It is also worth noting that different α-synuclein truncation forms may confer differences in α-synuclein functionality and speed of pathogenic progression (23). Our study indeed indicates stronger immune responses to the α-synuclein peptide, rather than the whole α-synuclein protein (**Fig. S2C**). It is also important to note that the model and successful disease penetrance is dependent on individual immune competency and different environmental elements and microbiome investigation should examine the way the whole protein is processed and presented by the MHCII machinery on Antigen-Presenting Cells, and whether border-associated macrophages mediate the neuroinflammation response as it has been demonstrated in an alpha-synuclein model of PD (24).

Misfolded α-synuclein can be detected in blood samples of PD patients by a seed amplification assay (SAA), and this is negatively correlated with disease duration, indicating that α-synuclein is a reliable predictor of PD onset in prodromal cases (5). It would be interesting to further assess if, together with α-synuclein, immune activation also represents a valid predictor of disease onset. Patients’ metadata analysis will be crucial to confirm whether CD4+ T cell activation is concomitant with elevated α-synuclein levels in humans in prodromal cases. Interestingly, both immune cells composition and α-synuclein levels vary in humans with aging (18), suggesting that age factor elevates the probability of CD4+/ α-synuclein interactions. While this work uncovered a causative link between a specific immune reaction to α-synuclein and the ontogenesis of PD-like features in mice, future investigation should also aim to decipher the specific mechanisms on how α-synuclein specific to CD4+ T cells travels to the brain, considering the Braak’s theory that demonstrates specific routes of α-synuclein deposition (25, 26). In that context, this study offers an ideal mouse model to study prodromal PD. There is no known cause or cure of PD, with 90% of cases identified as non-genetic or idiopathic with unknown derivations. Despite this, current models of PD in research are either transgenic or neurotoxic, imposing an intervention to replicate late-stage hallmarks of the disease. For example, 6-OHDA rodent experiments involve lesioning structures in the nigrostriatal pathway to induce Parkinsonian motor symptoms (27), whilst pre-formed fibril models inject α-synuclein aggregates directly to local regions in the brain to induce degeneration (28). While these models are advantageous in identifying existing intracellular and systemic mechanisms in PD and their relationship with symptoms, what is not known or modelled, is the cause of these events. We acknowledge that not all idiopathic PD cases are expected to have an immune origin. However, common processes are likely present, and the present α-synuclein immune-induced mouse model offers the opportunity to study the mechanisms underlying PD ontogenesis from molecular to organismal levels. This new model also enables monitoring prodromal, early stages of disease preceding symptom development, and investigating preventative measures for PD that aim to block α-synuclein (29) and/or inflammation in the brain(30).

## ACKNOWLEDGMENTS

Prof Glenda Halliday and Prof Riccardo Natoli for their expertise, and Prof Jana Vukovic for the Ly6b and CD3 antibodies. We thank Dr. Arti Medhavy for her flow cytometry expertise.

## AUTHOR CONTRIBUTIONS

Design of the experiments: RP, JP, CP, AB & ND. SG supplied the α-synuclein knockout mice. RP, JP, JM, IZ, ZX, RB & AR performed the experiments and RP, TX, JM & ZX performed the analysis. RP & ND prepared the figures and wrote the manuscript. AB edited the manuscript.

## STAR METHODS

### KEY RESOURCES TABLE

| REAGENT or RESOURCE | SOURCE | IDENTIFIER |
| --- | --- | --- |
| Antibodies |  |  |
| Tyrosine Hydroxylase Polyclonal Antibody | Thermo Fisher Scientific | Cat# PA1-4679; RRID: AB_561880 |
| GFAP Polyclonal Antibody | Thermo Fisher Scientific | Cat# PA5-85109; RRID: AB_2792257 |
| Anti Iba1, Rabbit antibody | FUJIFILM Wako Shibayagi | Cat# 019-19741; RRID: AB_839504 |
| Ly-6B.2 Monoclonal Antibody (7/4) | Thermo Fisher Scientific | Cat# MA5-16541; RRID: AB_2538046 |
| Anti-CD3 antibody [CD3-12] | Abcam | Cat# ab11089; RRID: AB_2889189 |
| Anti CD4: Rat IgG2a, k antibody | BD Biosciences | Cat# 553927; RRID: AB_395142 |
| Anti-A-synuclein filament antibody [MJFR-14-6-4-2] | Abcam | Cat# ab209538; RRID: AB_2714215 |
| Donkey Anti-Rabbit IgG (H+L) Polyclonal Antibody, Alexa Fluor 555 Conjugated | Molecular Probes | Cat# A-31572; RRID: AB_162543 |
| Donkey Anti-Rabbit IgG (H+L) Antibody, Alexa Fluor 488 Conjugated | Molecular Probes | Cat# A-21206; RRID: AB_2535792 |
| Donkey anti-Sheep IgG (H+L)<br>Secondary Antibody, Biotin | Thermo Fisher<br>Scientific | Cat# A16045, RRID: AB_2534719 |
| Cy2-AffiniPure Donkey Anti-Rat<br>IgG (H+L) (min X<br>Bov,Ck,Gt,GP,Sy<br>Hms,Hrs,Hu,Rb,Shp Sr Prot)<br>antibody | Jackson<br>ImmunoResearch<br>Labs | Cat# 712-225-150, RRID: AB_2340673 |
| Streptavidin Cy2 | Jackson<br>ImmunoResearch<br>Labs | Cat# 016-220-084, RRID: AB_2337246 |
| Streptavidin 647 | Jackson<br>ImmunoResearch<br>Labs | Cat# 016-600-084, RRID: AB_2341101 |
| Brilliant Violet 605(TM) anti-mouse/human CD45R/B220<br>antibody | BioLegend | Cat# 103243, RRID: AB_11203907 |
| APC anti-mouse/human CD11b<br>antibody | BioLegend | Cat# 101212, RRID: AB_312795 |
| BUV395 Rat Anti-Mouse Ly-6G<br>antibody | BD Biosciences | Cat# 563978, RRID: AB_2716852 |
| Brilliant Violet 421(TM) anti-mouse CD3epsilon antibody | BioLegend | Cat# 100335, RRID: AB_10898314 |
| PerCP/Cyanine5.5 anti-mouse<br>CD3 antibody | BioLegend | Cat# 100218, RRID: AB_1595492 |
| Alexa Fluor(R) 700 anti-mouse<br>CD4 antibody | BioLegend | Cat# 100536, RRID: AB_493701 |
| BV605 Rat Anti-Mouse CD4<br>Antibody | BioLegend | Cat# 563151, RRID: AB_2687549 |
| BUV395 Rat Anti-Mouse CD8a<br>antibody | BD Biosciences | Cat# 563786, RRID: AB_2732919 |
| PE/Cyanine7 anti-mouse CD11c<br>antibody | BioLegend | Cat# 117318, RRID: AB_493568 |
| PerCP/Cyanine5.5 anti-mouse Ly-6C antibody | BioLegend | Cat# 128012, RRID: AB_1659241 |
| 7-Aminoactinomycin D | Invitrogen | Cat# A1310 |
| Pacific Blue(TM) anti-mouse/human CD44 antibody | BioLegend | Cat# 103020, RRID: AB_493683 |
| Alexa Fluor(R) 700 anti-mouse<br>IL-17A antibody | BioLegend | Cat# 506914, RRID: AB_536016 |
| IFN gamma Monoclonal Antibody<br>(XMG1.2), APC, eBioscience | Thermo Fisher<br>Scientific | Cat# 17-7311-82, RRID: AB_469504 |
| Alexa Fluor(R) 488 anti-mouse<br>FOXP3 antibody | BioLegend | Cat# 126406, RRID: AB_1089113 |
| PE anti-mouse GM-CSF antibody | BioLegend | Cat# 505406, RRID: AB_315382 |
| IL-13 Monoclonal Antibody<br>(eBio13A), PE-Cyanine7,<br>eBioscience | Thermo Fisher<br>Scientific | Cat# 25-7133-82, RRID: AB_2573530 |
| Purified anti-mouse CD16/32 | BioLegend | Cat# 101301<br>can't find clone 2.4G2 |
| InVivoMAb anti-mouse CD28 | BioXCell | Cat# BE0015-1<br>Clone: 37.51 |
| Recombinant Murine IL-2 | PeproTech | Cat# 212-12 |
| Chemicals, peptides, and recombinant proteins |  |  |
| mouse α-synuclein protein | | |
| EndoClear™ human recombinant α-synuclein protein | AnaSpec | Cat# as-55555; Accession # NP_000336 |
| $\alpha$ -syn61-75 (EQVTNVGGAVVTGVT) | TC Peptide Labs | N/A |
| $\alpha$ -syn61-75 (EQVTNVGGAVVTGVT) | BioMatik | Cat# SP0001 |
| Pertussis Toxin from B. pertussis, Lyophilized (Salt-Free) | List Labs | Cat# 181 |
| Incomplete Freund Adjuvant (CFA) | Sigma Aldrich | Cat# F5506 |
| Complete Freund Adjuvant (CFA) | Sigma Aldrich | Cat# F5881 |
| M. tuberculosis H37Ra | Becton Dickinson | Cat# 231141 |
| V-52D DILUENT | American Veterinary Supply Co | N/A |
| Phosphate buffered saline powder, pH 7.4 | Sigma-Aldrich | Cat# P38135 |
| Paraformaldehyde 4% | Sigma-Aldrich | Cat# P6148 |
| Triton-X100 | Sigma-Aldrich | Cat# T8787 |
| Normal Donkey Serum | Merck | Cat# S30 |
| Bovine Serum Albumin | Sigma-Aldrich | Cat# A7906 |
| MOWIOL POLYVINYL ALCOHOL | Sigma-Aldrich | Cat# 81381 |
| Gelatine | VWR Chemicals | Cat# 24360.233 |
| Glycerol | Sigma-Aldrich | Cat# G5516 |
| Ethylene Glycol | Sigma-Aldrich | Cat# 324558 |
| Bovine serum albumin | Sigma-Aldrich | Cat# A7906 |
| DAPI 4',6-diamidino-2-phenylindole | Sigma-Aldrich | Cat# D9542 |
| Levodopa | Sigma-Aldrich | Cat# PHR1271 |
| S-(-)-Carbidopa | Sigma-Aldrich | Cat# C1335 |
| L-Ascorbic acid | Sigma-Aldrich | Cat# A4544 |
| DABCO 1,4 Diazabicyclo[2.2.2]octane | Sigma-Aldrich | Cat# D27802 |
| Ammonium chloride, 98+%, Thermo Scientific Chemicals | ThermoFisher Scientific | Cat# 123340250 |
| Potassium Bicarbonate | ThermoFisher Scientific | Cat# A12429.36 |
| Ethylenediaminetetraacetic acid (EDTA) (disodium salt) | Bio-Strategy Ltd |  |
| Iscove's Modified Dulbecco's Medium (IMDM) | ThermoFisher Scientific | Cat# 12440053 |
| Phorbol 12-Myristate 13-Acetate (PMA) | Sigma-Aldrich | Cat# P8139 |
| GolgiStop Protein Transport Inhibitor (containing monesin) | BD Biosciences | Cat# 554724 |
| Ionomycin | Sigma-Aldrich | Cat# I0634 |
| Heat Inactivated Fetal Calf Serum (HI FCS) | Sigma-Aldrich | Cat# F9665 |
| B2 Mercaptoethanol | ThermoFisher Scientific | Cat# 21985023 |
| OneComp eBeads | ThermoFisher Scientific | Cat# 01-1111-41 |
| Trypan Blue Solution | Sigma-Aldrich | Cat# T8154 |
| Critical commercial assays |  |  |
| eBioscience™ Foxp3 / Transcription Factor Staining Buffer Set | ThermoFisher Scientific | Cat# 00-5523-00 |
| Deposited data |  |  |
| Raw and analyzed data | This paper |  |
| Code | This paper | Github: |
| Experimental models: Organisms/strains |  |  |
| Mouse: C57BL/6J | Charles River Laboratory | Strain # RRID: |
| Mouse: C57BL/6J | The Jackson Laboratory | Strain #000664; RRID: IMSR_JAX:000664 |
| Mouse: B6.129S7-Rag1tm1Mom/J | The Jackson Laboratory | Strain #002216; RRID: IMSR_JAX:002216 |
| Mouse: α-Syn<sup>-/-</sup> | | Strain # RRID: |
| Software and algorithms |  |  |
| Fiji | ImageJ | Version 2.0.0, RRID:SCR_002285, https://imagej.net/software/fiji/ |
| QuPath | N/A | N/A |
| Prism v9.0 | GraphPad | Version 9.0, RRID:SCR_002798, https://www.graphpad.com/scientific-software/prism/ |
| FACSDiva | BD Biosciences, NSW, Australia | N/A |
| FlowJo v10.7 | BD Biosciences, NSW, Australia | N/A |
| DVC Analytics | Techniplast | N/A |
| DigiGait (version 16.A.2) | Mouse Specifics, Inc. | N/A |
| EthoVision XT | Noldus Technology | N/A |
| Matlab | MathWorks | N/A |
| Python | N/A | N/A |

#### Mice

All animal procedures were conducted in accordance with national and international guidelines and approved by the institutional animal committee of The Australian National University (Ethics protocol #2018/55 & 2021/49) and The University of Queensland (Ethics protocol #2022/AE000817). Mice from two different laboratories, Charles River Laboratory and the Jackson Laboratory, Australian BioResources Australia, were utilized in this study. No difference in blood immunophenotyping and cell counting were observed in these different WT mouse strains. 2 months old (∼20g) mice were housed in groups of maximum five in standard laboratory cages under controlled conditions (12 h light/dark cycle, 22±2°C temperature, 50±5% humidity) and provided with standard rodent chow and water ad libitum. Males and females were both used, and all mice were acclimated to the experimental facility for at least one week before the initiation of any experimental procedures. We did not observe any difference in immune induction and neurological and behavioural symptoms in males and females. To minimize potential confounding factors related to different housing conditions, different experimental groups (*i.e.* non injected, sham and α-synuclein protein or α-syn_61-75_ injected mice) were housed in the same cage. The mice were monitored bi-weekly, finding no evidence of changes in health or behavior between non-injected mice and mice only receiving adjuvants (*i.e.*, sham; data not shown).

#### α-synuclein immunization

Based off the Experimental Autoimmune Encephalomyelitis (EAE) model, mice aged 8 –10 weeks, unless otherwise specified, and minimum 20 g were immunized subcutaneously with 150 μg α-synuclein (AnaSpec, TC Peptide Labs, or BioMatik) emulsified 1:1 (v/v) with Complete Freud’s Adjuvant (CFA, Sigma-Aldrich) and heat inactivated Myobacterium Tuberculosis (MT, Becton Dickinson; 4 mg/mL in CFA). Injected variants of α-synuclein included Mouse Recombinant protein, Human Recombinant protein, or custom-synthesised sequence α-syn15aaEQVTNVGGAVVTGVT, referred as α-syn_61-75_. The latter was found as an epitope to CD4+T cell specific responses in preclinical Parkinson’s patients (Arlehamn et al., 2020). We performed a total of 5 batches for α-synuclein protein and 10 for α-syn_61-75_ peptide. Sham mice received CFA-MT and pertussis toxin alone. Mice were similarly weighed and clinically monitored biweekly according to standard EAE scoring procedures (data not shown).

Emulsifications were prepared in TissueLyser II, or Minilys Emulsion PowerKit using 30Hz oscillation frequency over a minimum of 15 minutes. On days 0 and 2, 300 ng of pertussis toxin was administered intraperitoneally to all immunized mice.

#### α-synuclein immunization booster

To determine immune memory of the α-synuclein immunization, we performed booster injections in two different batches (males and females) which displayed the same results. As customary to standard immunization procedures, booster injections of the α-syn_61-75_ were repeated halfway through the disease course (*i.e.,* 4 weeks post immunization, 4 wpi) at the same dose and concentration in Incomplete Freud’s Adjuvant (IFA, Sigma-Aldrich) that lacks bacillus. Mice did not receive additional injections of pertussis toxin to prevent hypersensitivity reactions. Sham mice received IFA alone. These mice were additionally bled one week after receiving booster injections.

#### Immune cell isolation and flow cytometry analyses

##### Blood collection

Blood was collected by retro-orbital sinus, superficial temporal vein, or tail vein prior to 1-2 weeks before inoculation (baseline), 1, 5 and/or 8 wpi; or cardiac puncture (terminal 8 weeks only).

##### Lymph node preparation and in vitro culture

Cervical lymph nodes were mashed through 70 μm nylon filters and rinsed with 2% FCS (Sigma-Aldrich) in PBS. Cells were centrifuged and resuspended in 2% FCS in PBS at 10^7^ live cells/ mL. Viable cell numbers were determined by trypan blue exclusion. Cells were resuspended in complete Iscove’s Modified Dulbecco’s Medium (IMDM, ThermoFisher Scientific) containing 50 nM Phorbol 12-Myristate 13-Acetate (PMA, Sigma-Aldrich), 750 nM Ionomycin (Sigma-Aldrich), 0.1% v/v GolgiStop (BDBiosciences), αCD28 (1:250, BioXCell) and IL-2 (1:1000, PeproTech) and plated 10^6^ cells/ well. Cells were incubated at 37°C 5% CO2 for 4 h in 5 μg/mL α-syn61-75, 125μg/mL α-syn, or CIMDM alone.

##### Haematology

Whole blood was processed 1:1 (v/v) in PBS via ADVIA2120i or 1:25 (v/v) in Diluent (see key resources table) via MindRay BC-5000Vet haematology analyzers. Haematological parameters include absolute counts of white blood cells; red blood cells; platelets; haemoglobin; neutrophils; monocytes; lymphocytes; eosinophils; basophils; and solid blood volume percentage, also referred as haematocrits, defined by number of RBCs and MCVs as a percentage of total blood volume. We observed diversity of immune cells in each sample and therefore used difference between baseline and 8 wpi to consider these intrinsic differences.

##### Extra and Intracellular Staining

Cells collected from blood or lymph node culture were lysed with red blood cell lysis buffer (150 mM Ammonium Chloride, ThermoFisher Scientific; 10 mM Potassium Bicarbonate, ThermoFisher Scientific; and 0.1 mM NaEDTA, Bio-Strategy Ltd) 3 times for 3 min, incubated in CD16/32 (BioLegend) for 10 min, and stained for surface markers for 30 min. Washing in 2% FCS in PBS was performed between each step. For intracellular cytokine staining, cells were stimulated for 6 h with 7.37 nM PMA and 500 nM Ionomycin in complete IMDM (CIMDM), prepared by adding 0.05% (v/v) PMA dilution and 0.05% Ionomycin (v/v) to 10 mL of complete IMDM. 0.01 (v/v) GolgiStop was added in the last 4 h of stimulation. Stimulated cells were washed with PBS and incubated with 7-Aminoactinomycin D (1:1000; Invitrogen) for 20 min. Cells were washed in 2% FCS in PBS and then fixed, permeabilized, and stained with intracellular cytokines using the materials and protocol included in the FoxP3 staining kit (eBioscience).

##### Flow Cytometry and Analysis

Flow cytometry was run on a BD Fortessa (BD Biosciences) or LSRII flow cytometer (BD Biosciences), data recorded on BD FACS Diva, and analyzed on FlowJo software. All gating strategies are shown in Fig S.2 and referred to in the appropriate figure legends. Large datasets of flow cytometry populations from blood samples were further processed. Data was normalised and homogenised with a GPU accelerated implementation of Harmony using batch and time point. Subsequently, populations were identified with UMAP segments run on all cells and verified with 2x concurrent UMAPs and X2 concurrent PacMAP dimensionality analysis. The top 5 features derived from all cell population frequency and MFIs were then compared between groups using Boruta feature extraction run 5 times with 3-fold cross validation, accumulating feature importances. These features were then combined into a generalised linear model (binomial distribution) for discrimination between sham and α-syn_61-75_ classes to explore how these factors interacted with each other. (ref github)

#### Histology

##### Brain collection

Mice were anesthetized with isoflurane and transcardially perfused with 4% paraformaldehyde (PFA). Brains were carefully removed, post-fixed in 4% PFA overnight, and washed in PBS. Sagittal or coronal brain sections (40 μm thick) were cut using a cryostat (Leica CM3050 S) and slices were stored for long term storage at –20 degrees celsius in ethylene glycol.

##### Immunohistochemistry

Tissue sections were washed in PBS and permeabilized with Triton X-100 twice each for 5 min. Non-specific binding sites were blocked for 2 h in blocking solution (10% (v/v) normal donkey serum, 0.25% (v/v) Triton X-100, 2% (v/v) bovine serum albumin in 0.01M PBS). Sections were then incubated overnight at 4°C in primary antibodies, and the following day, incubated for 2-3 h in secondary antibodies and 1 h in tertiary antibodies in blocking solution. Sections were washed in PBS 3 times for 15 min between each step. Sections were counterstained with 5 μM DAPI for 10 min, washed in PBS 4 times for 5 min, mounted on glass slides with gelatine, and coverslipped with Mowiol mounting medium.

##### Image acquisition

Image acquisition was conducted on A1 Nikon Confocal, Zeiss LSM 510 META Confocal, Leica TCS SP5, MetaSystems Metafer VSlide Scanner, and Diskovery Spinning Disk Confocal. Images were taken of the striatum (at 4x or 10x) and substantia nigra (at 20x) on sagittal planes 0.96 mm - 1.56 mm lateral from midline (Paxinos atlas). Images were stitched as required to obtain whole structures. MJFR −14-6-4-2+ cells were imaged at 63x in 2-3 locations of the substantia nigra pars compacta per slice. Lateral ventricle images were taken at 20x on coronal planes, approximately –1 mm relative to Bregma. The same confocal microscope and parameters for offset, pinhole, gain and laser intensity were kept constant within batches and experiments.

##### Image analysis

Maximum intensity was taken from z-stack images, and each analysis was averaged over 2-4 slices per mouse. Cell counting was performed manually on Fiji Image J except for NeuN+ cells, which were calculated by positive cell detection on QuPath (link macro). Cell counts were taken as a density of the region of interest’s area. On Fiji Image J, mean immunofluorescent intensity was calculated as a ratio of background signal, and neuropil involved taking the signal area percentage (link macro). CD4+ cell density was calculated on QuPath by the number of DAPI cells positive for CD4+ signal exceeding a single threshold of 1000, within a 5-uM radius from the nuclei. Density was calculated over a consistent area of the lateral ventricle containing the choroid plexus. A similar method was used for MJFR −14- 6-4-2+ cell count and fluorescent intensity when colocalised with Th+ cells (link macro).

#### Behaviour

To assess and monitor locomotor activity mice were recorded at baseline (pre-induction), and a minimum of 4 and 8 wpi. Mice were acclimatized to the testing room for 30 minutes and weighed prior to testing. Behavioural recordings were performed by the same experimenter and kept at a consistent LUX and time of day between experiments within batch.

##### Digitally Ventilated Cages (DVC)

At 100 days wpi, C57Bl6JAusb females (no injected, sham and α-synuclein, immunized mice) were single housed in Digitally ventilated cages (DVCs) for 12 days with ad libitum access to food and water. Food was weighed before and after the 12 days, taking the difference in weights as a measure of food consumed. Food consumption was then taken as a ratio of individual weight difference over the 12 days. Each cage measured 30 cm x 18 cm x 13 cm and equipped with a digital ventilation system and movement sensors. Data from these sensors were logged digitally every 15 minutes for real-time monitoring and subsequent analysis on DVC® Analytics. Correlations between environmental parameters and behavioral or physiological responses of the mice were examined, such as movement and speed (data not shown), and Regularity Disruption Index (RDI). These values were recorded and taken as an accumulation of the 12 days. We show that RDI is a robust measure capable of detecting home cage activity patterns that could be related to rest/sleep-related disturbances during the disease progression.

##### Open Field

To assess general locomotor activity, mice were allowed to move freely for 1 to 5 min over 2 trials in an open field arena at 8 wpi. Initial batches were recorded in a 38.5cm diameter circular arena analysed on MATLAB (GitHub; revised from Zhang et al., 2020). Later testings were conducted in a square (50 cm x 50 cm x 40 cm) arena and analysed via EthoVision XT tracking software (Noldus Information Technology).

##### Rotarod

As a measure of coordination and balance control, mice underwent rotarod testing (Ugo Basile; 47600, IITC Life Science; 755 & series 8 software, and Panlab Harvard Apparatus; LE8200 (76–0237)). Mice were first trained on the rotarod at a constant speed of 10 rpm over a maximum of 180 seconds for 2 trials, 24 h prior to testing. Mice were gently placed back on the rod if they fell during training. Two different test protocols were conducted with 2 to 3 trials each, wherein mice were subjected to 10 to 40 rpm acceleration over 40 or 180 seconds. The time each mouse remained on the rotating rod was recorded as the latency to fall or perform a full rotation. Results were consistent between the 2 test protocols (Fig S4.X).

##### Grip Strength

Grip strength was quantified by testing the force exerted via the mouse’s grip on the T-bar of a force metre (Able Scientific; GT-3, and IMADA; DS2 Digital Force Gauge) when pulling from the tail. Recordings were averaged over 2 to 3 trials and taken as a ratio of the weight of the mouse at time of testing.

##### DigiGait

Video recordings were taken from the ventral view of mice running 20cm/s on DigiGait apparatus. Mice were individually tested on the treadmill, capturing between 3-10 sec of footage per trial, for 2 trials. Videos were analyzed on the corresponding DigiGait software, giving an output of around 30 metrics of mobility and gait kinematics. Data was processed using Principal Component Analysis (PCA) to extract the underlying structure and reduce dimensionality, facilitating interpretation of the dataset (link code).

##### Levodopa Treatment

α-syn_61-75_ immunized mice were treated with Levodopa-carbidopa for 15 consecutive days from 8 wpi. Each day 2.5 ug carbidopa/ g mouse (20 mM in DMSO) was administered by intraperitoneal injection, followed by single intraperitoneal injection of 10 ug levodopa/ g mouse (Ascorbic acid, and 3 mg/mL in H2O). α-syn_61-75_ controls received 100 uL saline over the 15 days.

#### Statistics

All data are presented as mean ± standard error of the mean (SEM) unless otherwise stated. Statistical analysis was performed using GraphPad Prism software (GraphPad Software Inc.). Differences between groups were assessed by one, or two-way analysis of variance (ANOVA) followed by Tukey’s post hoc test for multiple comparisons. A p-value of less than 0.05 was considered statistically significant.

## SUPPLEMENTAL INFORMATION

Supplementary Information is available for this manuscript. *Link

Correspondence and requests for materials should be addressed to. Supplementary material: Github, Matlab codes, Table PACMACs

## SUPPLEMENTARY FIGURE LEGENDS

**Figure S1. Immune cell activation by hematology in α-synuclein immunized mice.**

(A) Experimental groups. (B) Timeline including blood collection (red triangle), injection (purple). BL: Baseline, wpi: week post-injection. (C) Full blood counts of white blood cells (WBC), monocytes, neutrophils, eosinophils, and lymphocytes quantified as a ratio of count 1 wpi over baseline (pre-immunization). Two-way ANOVA condition effect F(2,555) = 8.983, p = 0.0002. No-induction: n = 10; sham n = 30, and α-syn immunized n = 14 mice. (D) Absolute monocytes in non-induced (n = 6), sham (n = 10), murine α-syn (n = 7), and hµman α-syn (n = 9) immunized mice at 4 wpi; Kruskal-Wallis test p = 0.0017. Dunn’s multiple comparisons of no-induction compared to murine α-syn p = 0.0021 and hµman α-syn p = 0.0029. (E) Neutrophils count in non-induced (n = 6), sham (n = 10), murine α-syn (n = 7), and hµman α-syn (n = 9) immunized mice at 4 wpi. One-way ANOVA condition effect F(3,28) = 5.065, p = 0.0063. Tukey’s multiple comparisons of no-induction compared to murine α-syn p = 0.0375 and hµman α-syn p = 0.0038. (F) Absolute Red Blood Cells (RBC) at baseline and 1 wpi; non-induced (n = 10), sham (n = 24-25), α-syn immunized (n = 14-15) mice, sham vs α-syn p = X. (G) Haematocrits (HCT) count at baseline and 1 wpi; non-induced (n = 10), sham (n = 24-25), α-syn immunized (n = 14-15) mice, sham vs α-syn p = X. (H) Haemoglobin (HGB) count at baseline and 1 wpi; non-induced (n = 10), sham (n = 24-25), α-syn immunized (n = 14-15) mice. Sham vs α-syn p = X. (I) Longitudinal White Blood Cell (WBC) count in non-induced (n = 3-10), sham (n = 5-20) and α-syn immunized mice (n = 5-15). Mixed effects repeated measures interaction effect F(6,74) = 6.94, p < 0.0001. Tukey’s multiple comparison to no-induction vs sham p < 0.0001, and α-syn p < 0.0001 at 1 wpi, and sham p = 0.0022, and α-syn p < 0.0001 at 8 wpi. (J) Longitudinal monocytes count in non-induced (n = 3-10), sham (n = 5-20) and α-syn immunized mice (n = 5-15). Mixed effects repeated measures interaction effect F(6,118) = 4.406, p = 0.0005. Tukey’s multiple comparison to no-induction vs sham p < 0.0001, and α-syn p < 0.0001 at 1 wpi; sham p = 0.0007, and α-syn p = 0.0353 at 8 wpi. (K) Longitudinal neutrophils count in non-induced (n = 4-10), sham (n = 5-20) and α-syn immunized mice (n = 4-15). Mixed effects repeated measures interaction effect F(6,74) = 7.43, p < 0.0001. Tukey’s multiple comparison to no-induction vs sham p < 0.0001, and α-syn p = 0.0014 at 1 wpi, and sham p = 0.0349 at 8 wpi. Data is shown as mean ± SEM. Significance represented with p < 0.05: *; p < 0.01: **; p < 0.001: ***, p < 0.0001: ****. Non-paired Student T test otherwise stated.

**Figure S2. Flow cytometry gating strategies and analysis.**

(A) Representative flow cytometry plots show the gating strategies in blood and lymph node cultures. After gating live single cells, CD3+B220-populations were gated for analysis of activated CD4 and CD8 populations, while CD3-CD19- were gated for analysis of CD11b, CD11c, and Ly6c populations. (B) Representative flow cytometry plots of GM-CSF and IL-17, after pre-gating CD4+CD44+ cells according to Fig S2. (C) IL-17+ and/or GMCSF+ populations as a percentage of CD4+ cells at baseline (pre-immunization) and 1 week post immunization (wpi) in non-induced (n = 3-5), sham (n = 4-10), and α-syn immunized (n = 3-10) mice. Mixed-effects repeated measures interaction effect F(4,30) = 3.470, p = 0.0191. Tukey’s multiple comparisons between α-syn and no-induction (p < 0.0001; 1 wpi & p = 0.0492; 8 wpi), and α-syn and sham (p = 0.0119; 1 wpi). Data points represent individual mice values. Data is shown as mean ± SEM. Significance represented with p < 0.05: *; p < 0.01: **.

**Figure S3. Cellular alterations in the α-synuclein immunized mice.**

No differences in immune cell populations in the substantia nigra (SN) at 8 wpi. including (A) CD3+ cell density in no-induction (n = 4), sham (n = 4) and α-syn_61-75_ (n = 5), One-way ANOVA F(2,10) = 0.1167, p = 0.8910, (B) CD4+cell density in no-induction (n = 5), sham (n = 10) and α-syn_61-75_ (n = 9), X, and (C) Ly6b+cell density in no-induction (n = 4), sham (n = 5) and α-syn_61-75_ (n = 5), Kruskal-Wallis test, p > 0.9999. D) No difference in dopaminergic cell (Tyrosine hydroxylase: Th) density in the substantia nigra pars compacta (SNc) at 8 wpi across controls, including female no-induction (n = 5), female sham (n = 11), and male sham (n = 9) mice, One-way ANOVA F(2,22) = 1.859, p = 0.1794. Sham mice pooled across sex for further analysis. E) Th+ density in the SNc at 8 wpi in sham (n = 20) and α-syn injected mice (n = 10), p = 0.0011. (F) SNc NeuN density in sham (n = 15) and α-syn injected mice (n = 5) at 8 wpi, p = 0.0421. (G) Th expression levels in the striatµm normalized to sham. Sham (n = 24) and α-syn injected mice (n = 14) at 8 wpi, p = 0.0383. (H) Th+ density in the VTA at 8 wpi. in sham (n = 5) and α-syn_61-75_ injected mice (n = 3), p = 0.6843. (I) IBA1+ percentage area in the SN of sham (n = 9) and α-syn immunized mice (n = 4) at 8 wpi, p = 0.0196, Mann-Whitney test. Data points represent individual mice values. Data is shown as mean ± SEM. Significance represented with p < 0.05: *; p < 0.01: **; p < 0.001: ***, p < 0.0001: ****. Non-paired Student T test otherwise stated.

**Figure S4. Behavioural deficits in the α-synuclein immunized mice.**

(A) Longitudinal weight measures of sham (n = 13), α-syn immunized (n = 5), and α-syn_61-75_ immunized (n = 12) mice. Two-way ANOVA interaction effect F(8,108) = 0.5539, p = 0.8132. (B) Food intake per weight gain over 12 days in no-induction (n = 4), sham (n = 4), and α-syn immunized mice (n = 6). One-way ANOVA connection effect F(2,11) = 2.121, p = 0.1664. No differences in controls between (C) rotarod test of non-injected (n = 5) vs sham mice (n = 30) at 8 weeks post immunization (wpi), p = 0.8794 and (D) grip strength test of non-injected (n = 5) vs sham mice (n = 29) at 8 wpi, p = 0.7449. (E) Longitudinal rotarod of sham (n = 13), α-syn immunized (n = 5), and α-syn_61-75_ immunized (n = 12) mice. Two-way ANOVA condition effect F(4,54) = 1.461, p = 0.2269. (F) Longitudinal grip strength for sham (n = 11-23), α-syn immunized (n = 8), and α-syn_61-75_ immunized (n = 14-19) mice. Two-way ANOVA interaction effect F(2,144) = 0.3059, p = 0.7369. (G) Left: Representative schematic of mouse movement tracking in open field arena from sham (top) and α-syn immunized mice (bottom). Middle: Distance travelled (m) in arena at 8 wpi in sham (n = 16) and α-syn immunized mice (n = 13), p < 0.0001, T-test with Welch correction. Right: Time spent at rest (s) at 8 wpi. in sham (n = 18) and α-syn immunized mice (n = 13), p < 0.0001, T-test with Welch correction. (H) Grip strength at 8 wpi in sham (n = 27) and α-syn mice (n = 20), p = 0.0181. (I) Accelerating rotarod test at 8 wpi in sham (n = 8) and α-syn mice (n = 5), p = 0.0139. (J) Density of DigiGait principal component analysis values between sham (n = 32) and α-syn_61-75_ immunized (n = 30) mice, p = 0.0178. (K) Regularity disruption index (RDI) in digitally ventilated cages cµmulative over 12 days from sham (n = 4) and α-syn immunized (n = 4) mice, F = X, One-way ANOVA. Data is shown as mean ± SEM. Significance represented with p < 0.05: *; p < 0.01: **; p < 0.001: ***, p < 0.0001: ****. Non-paired Student T test otherwise stated.

**Figure S5. Standard immunization of transgenic and booster mice does not affect the development of PD symptoms.**

(A) Whole white blood cells (WBC) of α-syn^-/-^ mice at baseline (pre-immunization; n = 11), compared to 1 week post immunization (wpi) α-syn^-/-^ sham (n = 9) and α-syn^-/-^ α-syn_61-75_ immunized (n = 9) mice. One-way ANOVA, F(2, 26) = 5.193, p = 0.0127. Dunnett’s multiple comparisons versus baseline: sham: p = 0.0161, α-syn^61-75^: p = 0.0267. (B) Dopaminergic cell (Tyrosine hydroxylase: TH) density in the substantia nigra pars compacta (SNc) in no-induction (n = 4) and sham (n = 10), p = 0.5778. (C) rotarod test of no-induction (n = 4) and sham (n = 10), p = 0.5104; and (D) grip strength of no-induction (n = 4) and sham (n = 10), p = 0.5950. (E) Absolute lymphocytes in wild type (WT) (n = 36) compared to Rag1^-/-^ (n = 15) mice at baseline, p < 0.0001, Mann-Whitney test. (F) Flow cytometry of CD4+ cells as percentage of live cells in WT (n = 16) compared to Rag1^-/-^ (n = 15) mice at baseline, p < 0.0001. (G) Whole white blood cells (WBC) of Rag1^-/-^ mice at baseline (n = 15), compared to 1 wpi Rag1^-/-^ sham (n = 5) and Rag1^-/-^ α-syn_61-75_ (n = 10). One-way ANOVA, F (2, 27) = 36.66, p < 0.0001. Dunnett’s multiple comparisons versus baseline: sham p < 0.0001, α-syn_61-75_ p < 0.0001. (H) Absolute white blood cells (WBC) quantified as a ratio of 8 wpi over baseline for no-induction (n = 8), sham (n = 13), α-syn_61-75_ (n = 8), sham booster (n = 7), and α-syn_61-75_ booster (n = 8). One-way ANOVA, F(4,39) = 28.18, p < 0.0001. Tukey’s multiple comparison of no-induction compared to sham (p = 0.0233) and α-syn_61-75_ (p = 0.0353); α-syn_61-75_ compared to boosted sham (p < 0.0001) and boosted α-syn_61-75_ (p < 0.0001). (I) Absolute neutrophils over time for no-induction (n = 8-10), sham (n = 13-20), α-syn_61-75_ (n = 13-16), sham booster (n = 11) and α-syn_61-75_ booster (n = 11). Two-way ANOVA condition effect F(4,208) = 13.08, p < 0.0001. Dotted line: booster injection at 4 wpi. Tukey’s multiple comparison post booster (5& 8 wpi) of α-syn_61-75_ vs α-syn_61-75_ booster p > 0.9999. (J) Flow cytometry of CD4+CD44+ percentage of live cells for sham (n = 3), α-syn_61-75_ (n = 5-9), sham booster (n = 11) and α-syn_61-75_ booster (n = 11). Two-way ANOVA condition effect F(3, 122) = 0.795, p = 0.4976. Tukey’s multiple comparison post booster of α-syn_61-75_ vs α-syn_61-75_ booster p = 0.9991. Data is shown as mean ± SEM. Significance represented with p < 0.05: *; p < 0.01: **; p < 0.001: ***, p < 0.0001: ****. Non-paired Student T test otherwise stated.

