## Supplementary figures and images for "Αlpha-Synuclein Induced Immune Response Triggers Parkinson’s Disease-Like Symptoms"

### Supplementary Figures

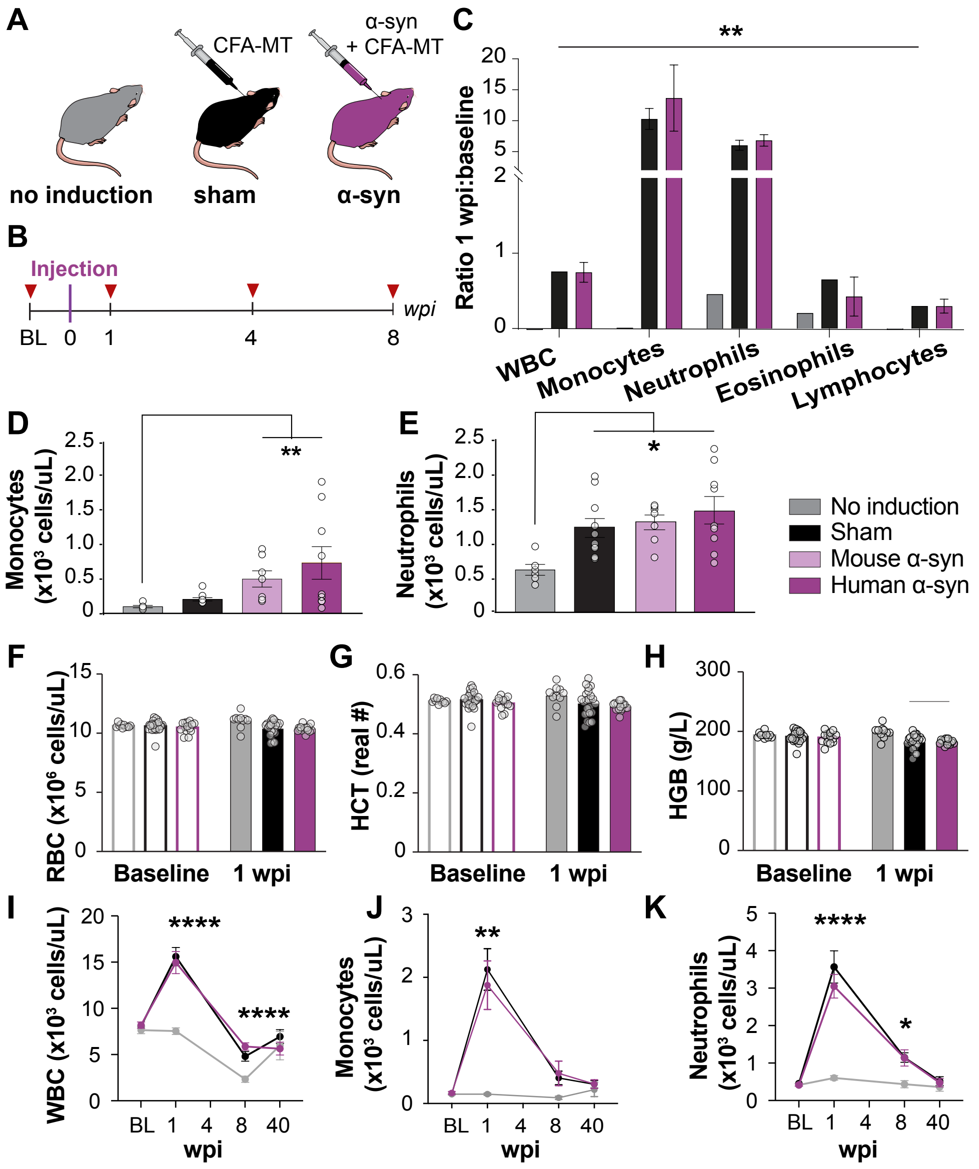

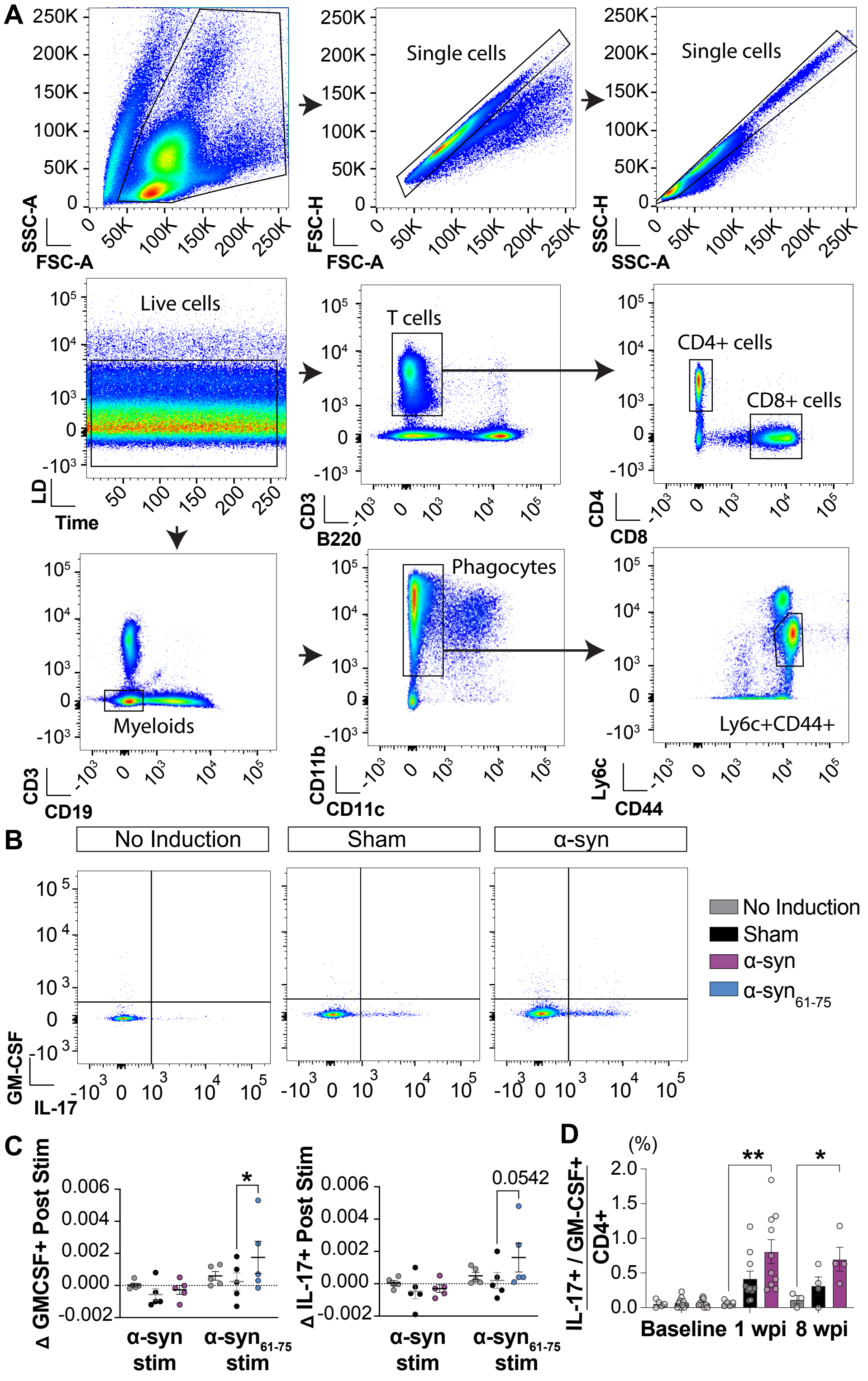

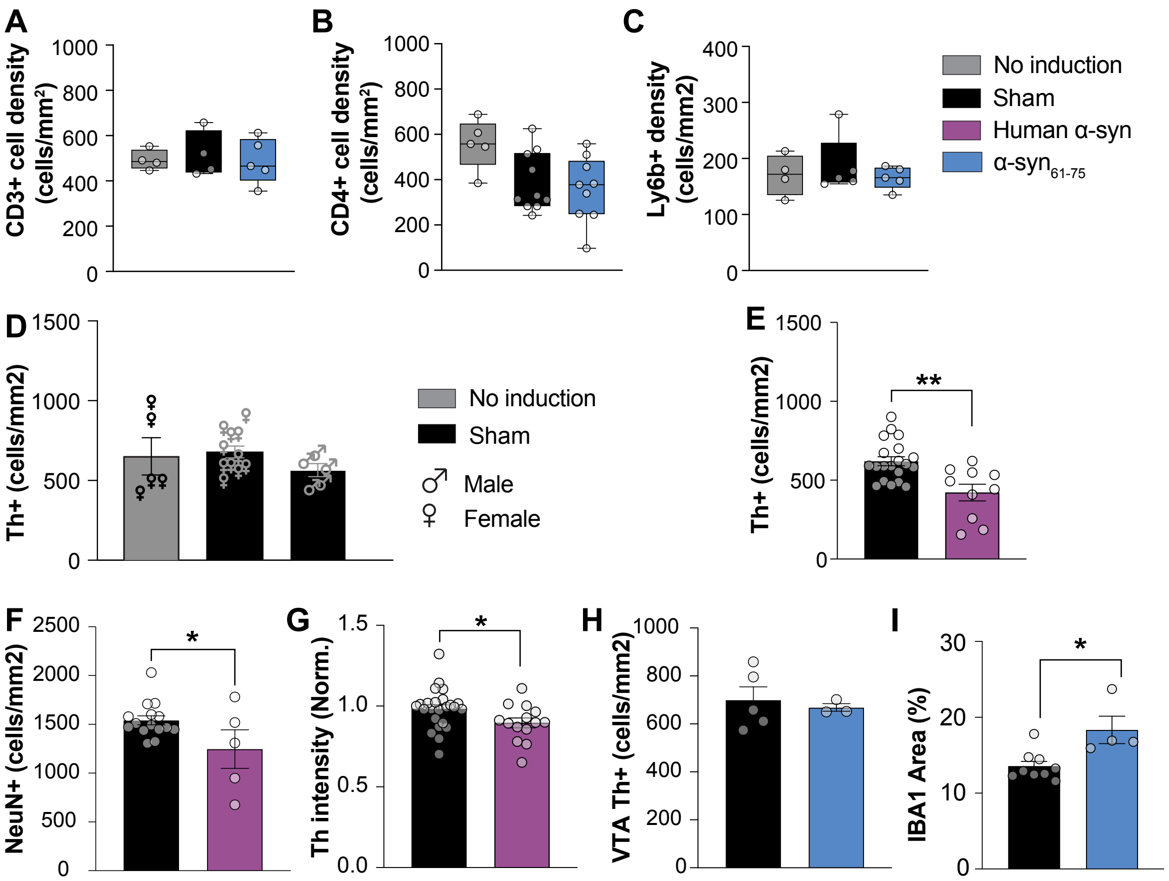

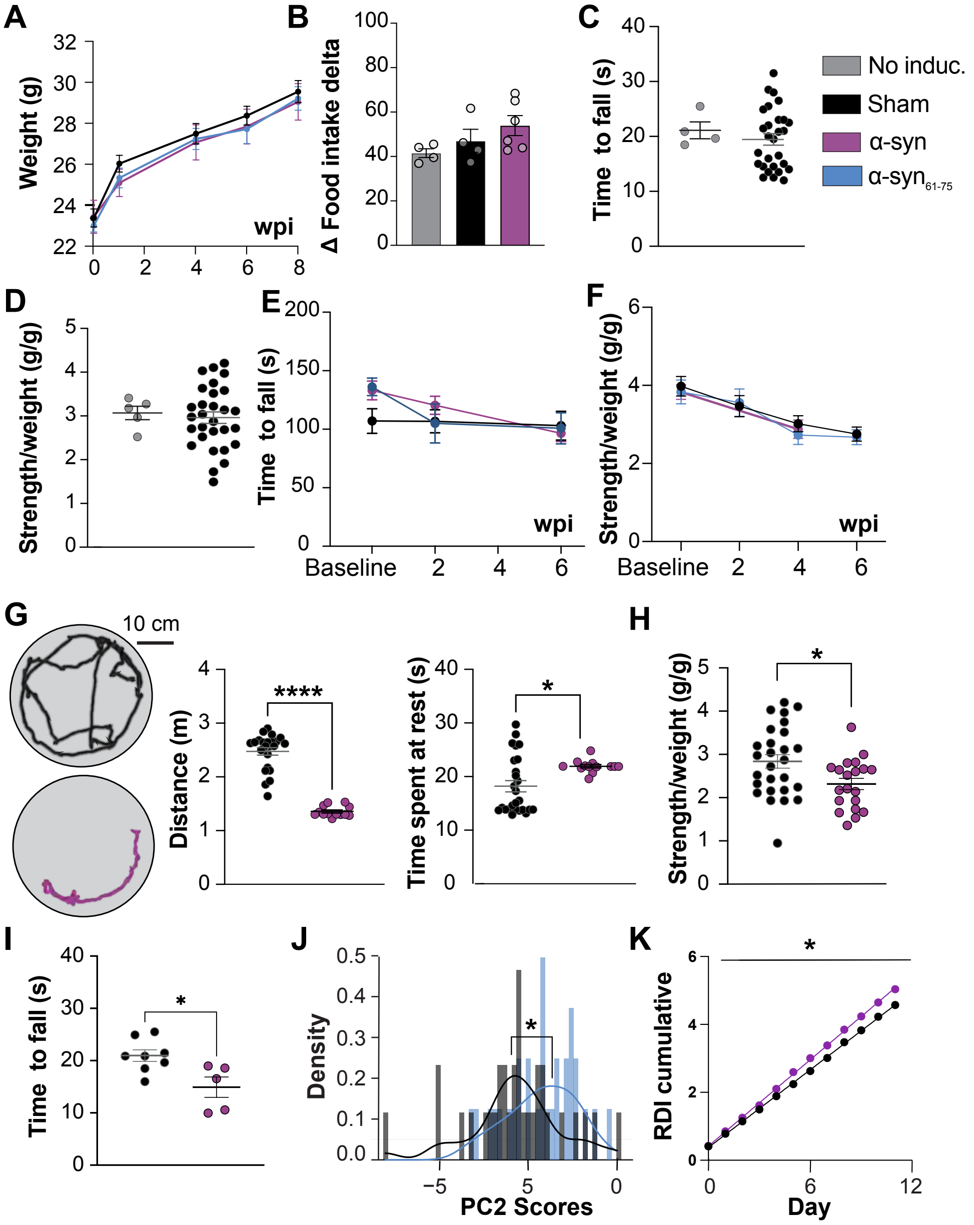

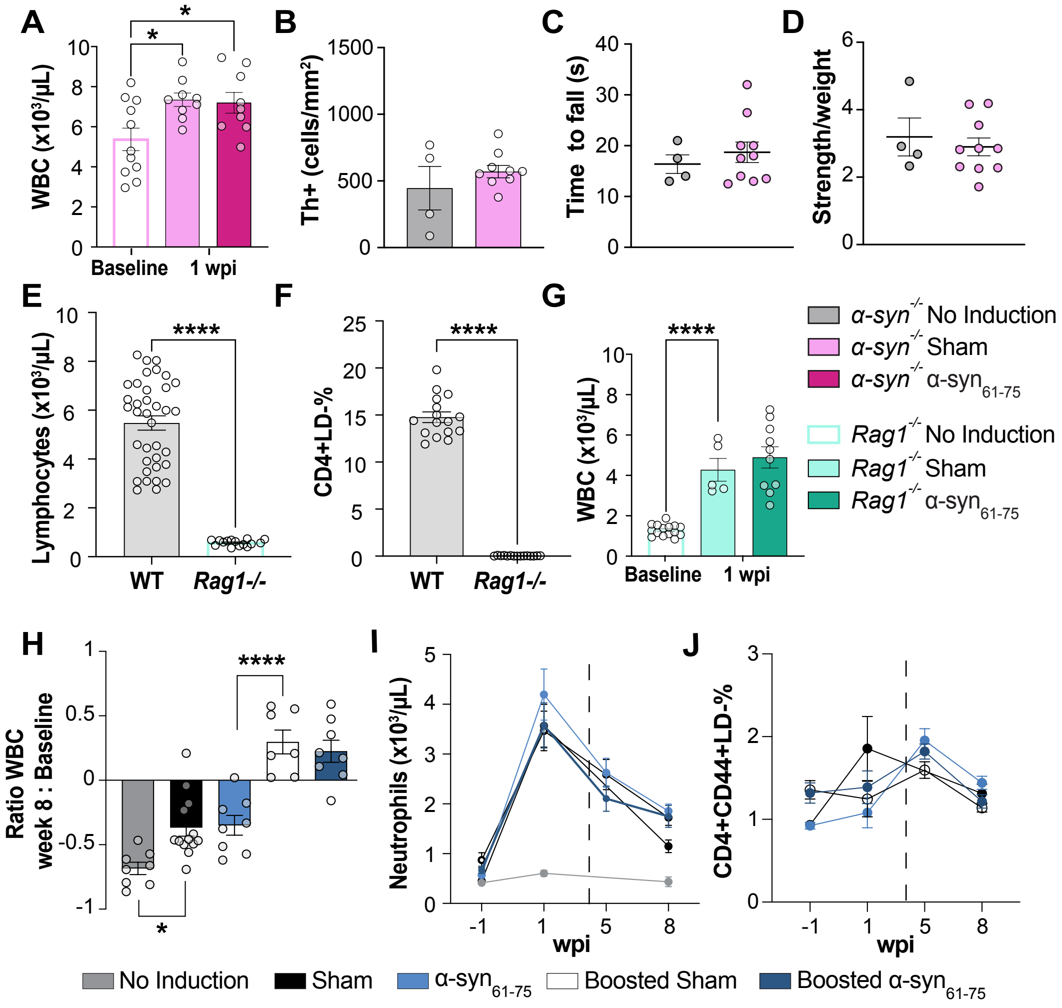

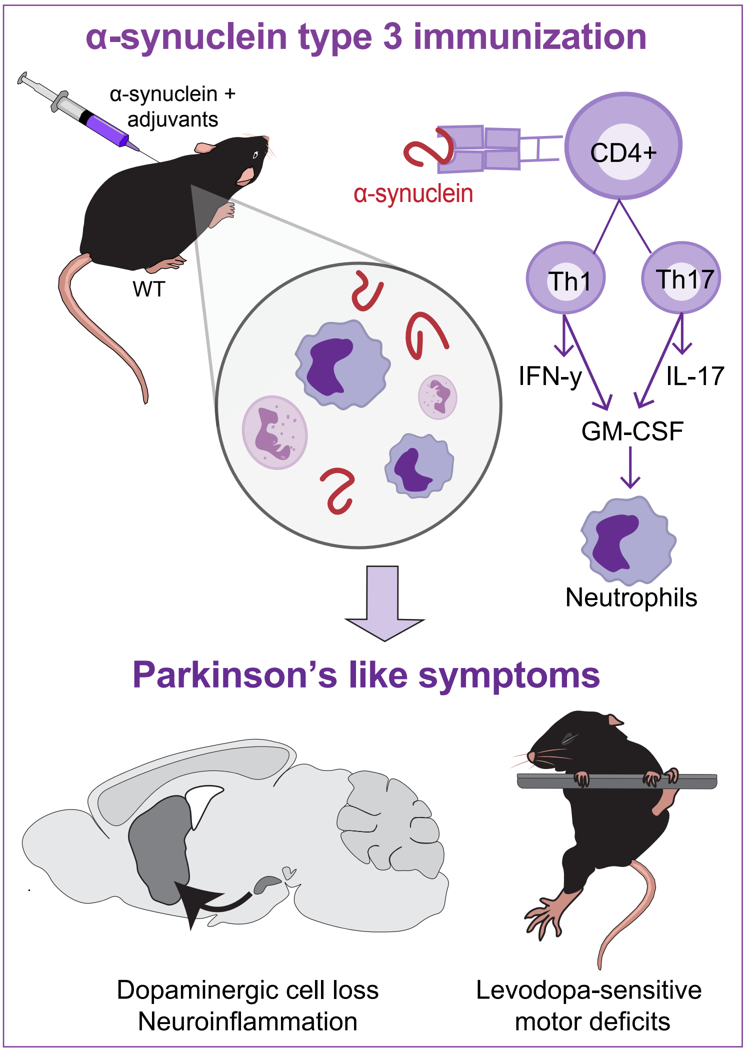
